# Ultraspecific somatic SNV and indel detection in single neurons using primary template-directed amplification

**DOI:** 10.1101/2021.04.30.442032

**Authors:** Lovelace J. Luquette, Michael B. Miller, Zinan Zhou, Craig L. Bohrson, Alon Galor, Michael A. Lodato, Charles Gawad, Jay West, Christopher A. Walsh, Peter J. Park

## Abstract

Primary template-directed amplification (PTA) is an improved amplification technique for single-cell DNA sequencing. We generated whole-genome analysis of 76 single neurons and developed SCAN2, a computational method to accurately identify both clonal and non-clonal somatic (i.e., limited to a single neuron) single nucleotide variants (SNVs) and small insertions and deletions (indels) using PTA data. Our analysis confirms an increase in non-clonal somatic mutation in single neurons with age, but revises estimates for the rate of this accumulation to be 15 SNVs per year. We also identify artifacts in other amplification methods. Most importantly, we show that somatic indels also increase by at least 2 indels per year per neuron and that indels may have a larger impact on gene function than somatic SNVs in human neurons.

## Introduction

Although somatic mutation has been studied extensively in cancer, investigation into the abundance, patterns, and effects of somatic mosaicism in non-neoplastic tissues has only recently begun^1–6^. Unlike tumor tissue in which somatic mutations of interest are shared by large clones, the majority of somatic mutations in normal tissues are typically shared by relatively few cells and are therefore difficult to detect. Recent studies have circumvented the technical difficulty of detecting rare somatic mutations by strategies including ultradeep sequencing of very small tissue samples^3,7^, exploiting naturally occurring genetically homogenous clones^8^, or clonal expansion of cells *in vitro*^5,9,10^.

Another strategy for detecting somatic mosaic mutations is to directly sequence DNA from a single cell. Single cell DNA sequencing (scDNA-seq) is capable of detecting the rarest somatic mutations (i.e., mutations private to a single cell) and can also provide information about cell lineage through shared somatic mutations^2,11^. This strategy is especially useful for examining somatic mutations in post-mitotic cells such as neurons, in which their presence is limited to single cells. A major bottleneck, however, has been the difficulty of amplifying the genome of a single cell accurately and evenly so that it can be sequenced by a high-throughput sequencer. For example, multiple displacement amplification (MDA)^12^, a popular amplification method for detecting point mutations, produces non-uniformity across the genome^13^ and often amplifies homologous alleles of diploid cells at different rates, leading to allelic imbalance^14^. These amplification artifacts pose substantial difficulties for identifying mutations from short-read sequencing data—especially mutations that are non-clonal and thus cannot be confirmed by sequencing multiple single cells. We previously used read-level phasing to filter artifacts in MDA samples and discovered an age-associated increase in somatic mutations in human neurons^6^, but were limited to analyzing mutations within a few hundred base pairs of germline SNPs (~15% of the genome). A newly developed single-cell amplification method called primary template-directed amplification (PTA) aims to reduce these artifacts by dampening the exponential nature of isothermal MDA^15^.

Here we compare single neurons amplified by both the MDA and PTA protocols from the prefrontal cortices of the same individuals and find that PTA substantially improves upon MDA. Nevertheless, conventional somatic SNV analysis (based on Genome Analysis Toolkit (GATK) best practices) of PTA data yields 0.9 false positives (FPs) per megabase, exceeding the mutation rate in some non-neoplastic cells by an order of magnitude^10^. We therefore developed SCAN2 (<u>S</u>ingle <u>C</u>ell <u>AN</u>alysis 2), a small mutation genotyper based on the SCAN-SNV^14^ model of allelic imbalance. SCAN2 detects non-clonal somatic SNVs and indels in scDNA-seq data with 60-fold fewer FPs per megabase than conventional calling and >5-fold fewer FPs than single-cell SNV genotypers. Somatic SNV detection in SCAN2 is greatly improved by a novel multi-sample approach that distinguishes mutations from artifacts based on 96-dimensional mutation signatures^16^; somatic indel calling is enabled by using multiple single cells to identify and remove sites with unusually high indel recurrences. SCAN2 confirms a previously reported signature of single nucleotide MDA artifacts^17^ and revises the rate of somatic SNV (sSNV) accumulation in aging neurons from the human prefrontal cortex^6^. Most notably, SCAN2 provides the first characterization of somatic indels in human neurons, revealing the yearly rate of indel accumulation and a bias toward genic regions. Two of four known clock-like indel signatures appear to be active in neurons; additionally, we find that aging-related neuronal indels are primarily enriched for indel signature 4 from the COSMIC catalog, a signature characterized by short deletions of 2-4 bp and with no known aetiology.

### PTA improves amplification quality and reduces artifact burden

The genomes of 25 single neurons from the prefrontal cortex (PFC) of eight neurotypical individuals were amplified by PTA and sequenced to 30-60X (**Fig. 1a**, **Supplementary Fig. 1a**, **Supplementary Table 1**). Compared to MDA-amplified single neuron WGS data from the same individuals^6^, PTA-amplified neurons showed several favorable characteristics, including substantial reduction in coverage variability across the genome (as measured by median absolute pairwise deviation (MAPD) and visual inspection of copy number profiles) and allelic imbalance, despite being sequenced to lower depth (**Fig. 1b-d**). Allelic balance measures the evenness of amplification between homologous alleles in a diploid cell; values near 0.5 indicate successful amplification of both alleles while values near 0 or 1 indicate loss of one allele. On average, only 37% of MDA-amplified genomes exhibited balance levels in the range of 0.3-0.7 compared to 68% of PTA genomes. We also found the rate of amplification failure among our PTA reactions to be low: only a single PTA neuron showed evidence of amplification failure in the form of near-complete loss of several haplotypes (**Supplementary Fig. 2**). However, we cannot rule out the possibility that *bona fide* mutations are the sources of these copy losses, meaning that none of the 25 PTA reactions failed. If these were indeed true mutations, then they are the only large-scale (>5 Mb, see Methods) copy number changes we detected in these neurons, which is unexpected given reports of pervasive copy number alterations in human neurons, especially from young individuals^19,20^.

**Figure 1.**
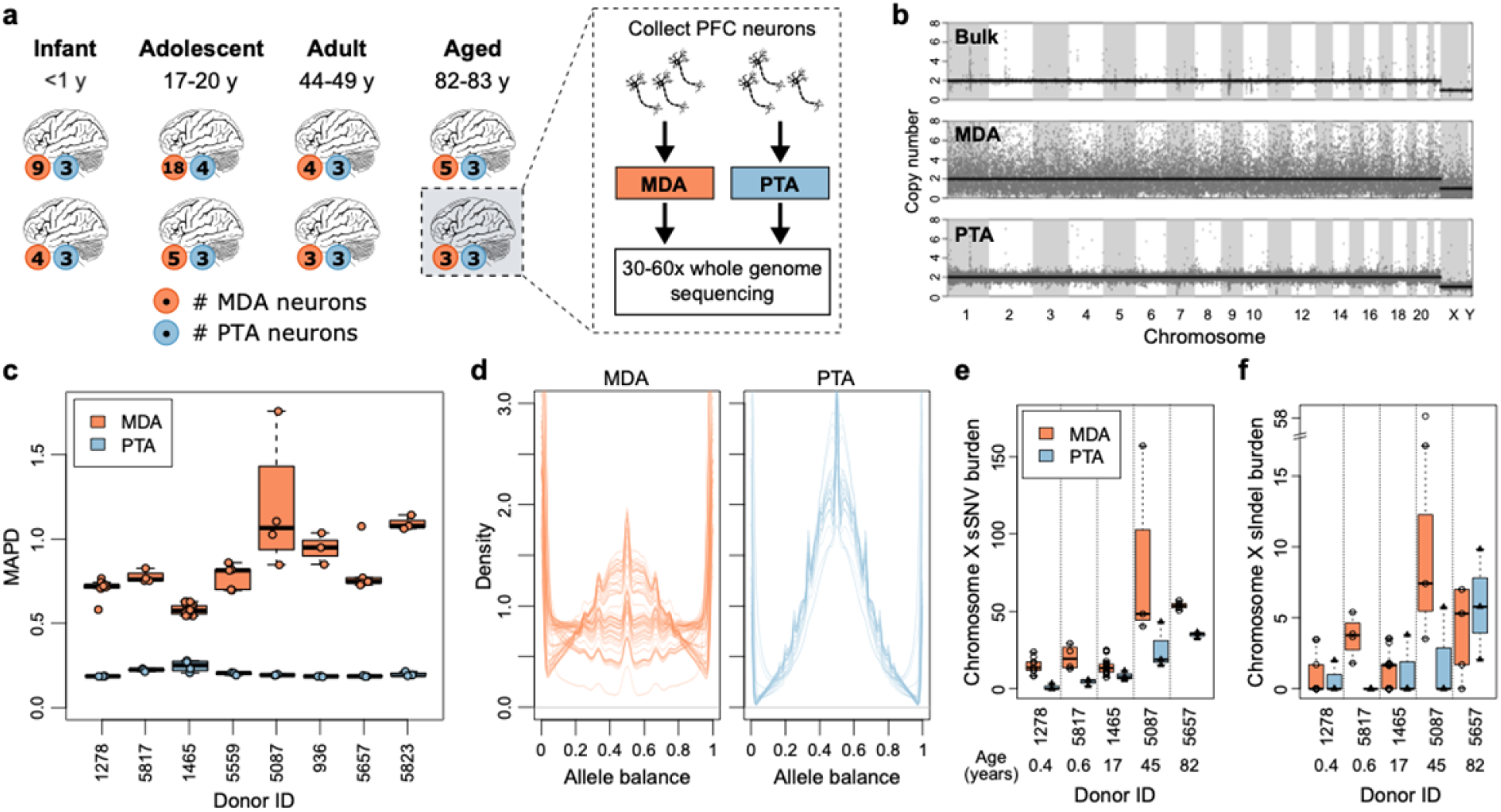
PTA improves over MDA at large and small scales. **a.** Study design. Single neurons were collected from the prefrontal cortices of brains of 8 individuals ranging in age from infantile to elderly. Single neurons were amplified by either PTA or MDA and then sequenced to high coverage. **b.** Representative copy number profiles for bulk (top), MDA-amplified (middle) and PTA-amplified (bottom) genomes. **c.** MAPD (median absolute pairwise deviation) for MDA-amplified and PTA-amplified neuronal genomes from the same individuals; lower values indicate better performance. The average MAPDs of MDA (0.75) and PTA (0.21) correspond to an average fluctuation in read depth between neighboring 50 kb windows of 68% and 14%, respectively. **d.** Allele balance for germline heterozygous SNPs in each sample. Each line corresponds to one single cell. Values near 0.5 indicate balanced amplification of homologous alleles; values near 0 or 1 indicate complete dropout of one allele. **e.** Sensitivity-adjusted somatic SNV (sSNV) burdens per X chromosome for 5 male individuals. **f.** Same as (e) for somatic indels (sIndels). Boxplot whiskers, furthest point at most 1.5× interquartile range.

Comparison of the numbers of somatic mutation calls between MDA and PTA amplified neurons from the same individual suggested specific types of artifacts introduced by MDA. In the absence of artifacts, the number of somatic calls should be similar in MDA and PTA from the same individual after correction for sensitivity, while a consistent excess of calls specific to one amplification method would indicate the presence of additional artifacts and allow estimation of the artifact rate. To measure the rates of high variant allele fraction (VAF) artifacts, we analyzed male X chromosomes since mutation detection in hemizygous regions is considerably less difficult than in diploid regions (Methods). MDA neurons displayed a median excess of 15.9 somatic SNVs and 3.7 somatic indels per haploid X chromosome, indicating that one should expect about 584 SNV and 136 indel high VAF artifacts per genome (**Fig. 1e-f, Supplementary Fig. 1b-c**). Notably, these MDA artifacts frequently occur with variant allele fractions (VAFs) of 100%, which is compatible with a previously proposed artifact model^21^ involving failure to amplify either the Watson or Crick strand of the initial DNA molecule. Artifacts caused by such single-stranded dropout do not leave the telltale signs of amplification artifacts (i.e., discordantly phased reads^21^ or improper VAFs^14^) and are often indistinguishable from true mutations.

### High specificity is critical for somatic mutation detection in healthy cells

The importance of single-stranded dropout MDA artifacts depends on how many mutations of interest exist in the cells being analyzed. For example, since human cells contain 3-4 million germline SNVs (>1000 SNVs/Mb), several hundred artifacts would have little effect on germline SNV discovery. Indeed, in the context of germline SNV detection, we estimate a false discovery rate (FDR) of <0.1% regardless of the amplification or analysis method (**Fig. 2a**). However, estimated FDR rates for MDA and conventional analysis of PTA are unacceptable when the mutations of interest are rare, as in somatic SNV detection in healthy single cells (0.1-1.0 sSNVs/Mb^5,6,9,10^). For MDA, we estimate a best-case scenario by assuming that the only FP errors are caused by single-stranded artifacts (see Methods). Under this assumption, we expect MDA FDRs of at least 17% (for cells with 1.0 sSNVs/Mb) to 68% (for cells with 0.1 sSNVs/MB); but in practice, higher MDA FDRs would be expected due to additional FPs from non-single-stranded artifacts. Although PTA produces fewer artifacts than MDA, single-cell-aware genotypers are critical for accurate sSNV calling in low mutation burden contexts: the conventional GATK best practices pipeline (with additional filtering) was recently estimated to produce 0.9 false positives (FPs) per megabase with ^~^80% sensitivity in PTA amplified cells^15^, corresponding to FDRs of 47% (1.0 sSNVs/MB) to 90% (0.1 sSNVs/MB) for typical healthy cells. In summary, both the optimistic MDA scenario and analysis of PTA by conventional genotypers are likely to produce unacceptable FDR levels in cells with low mutation burden. We therefore developed SCAN2, which achieves FDR < ^~^15% even for cells with very low mutation burden (0.1 sSNVs/Mb).

**Figure 2:**
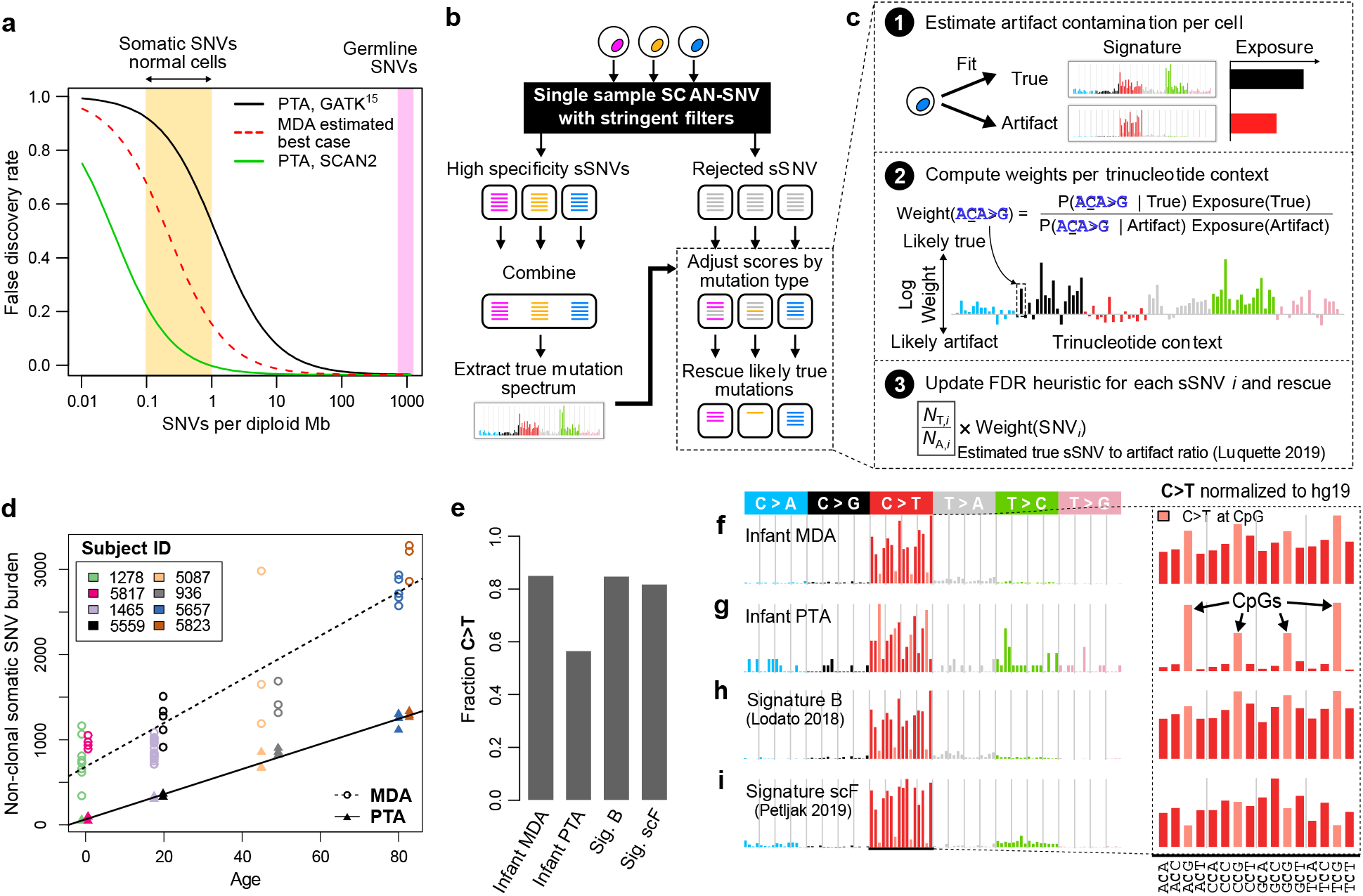
SCAN2 somatic SNV calling method and application to single human neurons. **a.** Estimated false discovery rates for MDA and PTA somatic SNV detection. The MDA best case scenario assumes that all single strand-dropout artifacts are erroneously called as true mutations. **b.** SCAN2 approach to multi-sample sSNV calling. SCAN2’s multi-sample approach is not phylogenetic and does not depend on sSNVs being shared by multiple single cells. It can therefore detect private mutations such as those in post-mitotic neurons. **c.** Candidate sSNVs are rescored separately for each single cell given the true mutation signature learned in panel (b). The likelihood of being generated by the true signature is computed for each mutation type and trinucleotide context (x-axis). This likelihood acts as a prior for a previously described heuristic that estimates the number of true mutations (*N_T,i_*) and artifacts (*N_A,i_*) with characteristics similar to the sSNV candidate *i*. **d.** Sensitivity-adjusted accumulation rate of somatic SNVs in PTA-(triangles) and MDA-(circles) amplified single human neurons. **e.** Fraction of C>Ts among sSNVs called by single sample SCAN2 in infant neurons and two previously published signatures. **f.** Mutational spectra of somatic SNVs called by SCAN2 in single-sample mode across 6 MDA neurons from 2 infants. Signature B is not subtracted from MDA calls. Right: rate of C>T mutations after normalizing by trinucleotide frequency in the human genome. **g.** Same as (f) for 6 PTA neurons from 2 infant donors. **h.** C>T rich neuron signature B reported in Lodato et al, 2018. **i.** MDA artifact signature scF reported by Petljak et al, 2019.

### SCAN2 accurately detects somatic SNVs and indels in PTA-amplified cells

SCAN2 is built on SCAN-SNV, a single-cell somatic SNV genotyper that accounts for allelic imbalance (the uneven amplification of homologous alleles)^14^. This is achieved by measuring the VAFs of heterozygous germline SNPs, which reflect the local allelic imbalance, near candidate sSNVs. SCAN2 incorporates two key advances over SCAN-SNV. First, we developed a novel multi-sample mutation signature-based approach to increase sensitivity for sSNVs and to provide a source of information orthogonal to VAF. In short, the method relies on differences between the mutation signatures of true somatic SNVs and amplification artifacts to rescue candidate sSNVs which are rejected by the SCAN-SNV model but are poor matches to the artifact signature. The approach operates in two passes (**Fig. 2b**, Methods): in the first pass, a set of high-specificity sSNVs is produced by running SCAN-SNV in single-sample mode with stringent calling parameters. These high-specificity sSNVs are then combined across cells to generate the mutation spectrum of the true mutational process. In the second pass, candidate sSNVs rejected in the first pass are re-assessed based on their mutation contexts and potentially rescued. To do this, exposures to the learned true mutation spectrum and a universal PTA artifact signature (for derivation of this signature, see Methods and **Supplementary Fig. 3**) are computed individually for each cell; then, based on the cell-specific mutation signature exposures, each mutation context is assigned a weight representing the likelihood of originating from the artifact signature; finally, the weights are used to adjust the SCAN-SNV FDR heuristic^14^ for rejected candidate sSNVs, allowing some candidates to be accepted (**Fig. 2c, Supplementary Fig. 4**). Although other multi-sample single-cell genotypers exist^22^, our method is unique in its capability to use cross-sample information to call private sSNVs, such as those that accumulate in post-mitotic cells.

The second key advance is the ability to call somatic indels in single-cell data. We hypothesized that, unlike artifactual sSNVs, artifactual indels are more likely to be recurrent owing to processes such as polymerase stutter^23^ and microhomology-mediated chimera formation^24^ that favor certain genomic regions. To identify indel artifacts, SCAN2 requires input from at least 2 distinct individuals to build a list of indel sites that are frequently mutated in multiple, unrelated cells. Candidate somatic indels are initially generated by a modified SCAN-SNV protocol and then screened against the multi-subject panel to remove recurrent candidates, as they are likely artifactual (Methods, **Supplementary Fig. 5**). While this filtration proves effective at removing many indel artifacts, it is expected to limit the ability to call somatic indels at hypermutable sites that are likely to occur in many individuals such as microsatellites^25^.

To assess the performance of SCAN2, synthetic diploid X chromosomes were simulated as previously described^14^. The multi-sample sSNV calling approach yields a mean sensitivity of 45.7%, 0.0143 FPs per megabase and mean FDR of 5.9% ± 6.8% at typical somatic mutation loads for healthy cells (**Supplementary Fig. 6a-c**). Notably, the multi-sample signature approach outperformed the single-sample approach in both sensitivity and FDR at every simulated mutation burden, ranging from 0.05 sSNVs/Mb-1.5 sSNVs/Mb. Furthermore, across the same mutation burden range, multi-sample SCAN2’s FDRs were lower than both Monovar^21^ and SCcaller^26^, two single-cell SNV genotypers developed for MDA-amplified single cells (**Supplementary Fig. 7**). We additionally found that SCAN2 is capable of accurately predicting the total mutation burden in PTA-amplified cells by estimating and correcting for detection sensitivity using germline SNPs (Methods, **Supplementary Fig. 6d**).

Assessment of somatic indel calling is complicated by the wide array of possible indels and the fact that indel detection sensitivity is affected by several indel characteristics, such as length and genomic context. We therefore generated a panel of indels with uniform representation across the ID83 classes, a set of 83 indel classes recently developed to enable mutation signature analysis of indels^27^, and used the synthetic diploid spike-in approach to score SCAN2’s sensitivity separately on each of the 83 channels. SCAN2 indel sensitivity ranged from 1.4%- 31%, with a clear pattern of reduced sensitivity for indels in tandem repeats greater than 4 units (**Supplementary Fig. 8**). Of particular interest, we found that cross-sample filtering considerably decreased sensitivity for single base insertions in long homopolymers, which are the primary constituents of two indel aging signatures in the COSMIC catalog (ID1 and ID2). We therefore expect that correcting for ID83 class-specific sensitivity will be crucial for somatic indel signature analysis. The FP rate for somatic indels did not exceed 0.001 FPs/Mb.

### Revised rates of nonclonal somatic SNVs in aging human neurons

SCAN2 identified 22,292 nonclonal sSNVs in the 51 MDA-amplified neurons using single sample calling and 7,174 across the 25 PTA neurons using the multi-sample approach informed by the PTA universal artifact signature. *De novo* signature extraction applied to the PTA sSNVs produced a single signature strongly resembling Signature A (cosine similarity 0.966), providing confirmation of the aging-associated signature we previously recovered from MDA-amplified neurons^6^ (**Supplementary Figure 9**). SCAN2 estimated the yearly rate of sSNV accumulation to be 14.7 sSNVs/year in PTA neurons compared to 25.7 sSNVs/year in MDA neurons from the same individuals. These rate estimates are not affected by differences in the multi-sample and single sample approaches, meaning that the difference is most likely explained by FP calls caused by greater MDA artifact burden (**Fig. 2d**) as was the case on the male X chromosomes. Nearly identical rates were produced by LiRA, a single-cell genotyper that uses an orthogonal approach both for calling sSNVs and for estimating the total sSNV burden per cell (**Supplementary Figure 10**). Importantly, although LiRA generates accurate calls based on read-level phasing, it is limited to genomic regions in close proximity to germline SNPs for phasing^21^; in contrast, SCAN2 can call mutations several kb from the nearest SNP and thereby generates a 5-fold increase in the number of sSNV calls.

To explore the nature of potential MDA artifacts, we focused on samples from the youngest subjects, infants, which should have the smallest true mutational burden. Amongst these samples, MDA neurons contain ^~^12-fold more SCAN2 sSNV calls than PTA neurons from the same individual after correcting for sensitivity, suggesting that infant MDA sSNVs can be regarded as a highly concentrated set of MDA artifacts. We first compared the infant MDA mutation spectrum with the higher quality infant PTA spectrum and found MDA sSNVs to be enriched for C>T mutations (85% vs. 59%, MDA vs. PTA) (**Fig. 2e-g**). Second, we noticed striking similarities between the infant MDA spectrum and two previously reported signatures that manifest in ways consistent with technical artifacts. Signature B (**Fig. 2h**) was previously reported in aging human neurons but did not increase with age^6^; Signature scF (**Fig. 2i**) was previously observed in MDA-amplified single cells but not in clonally expanded single cells from the same cell lines^17^. Third, we hypothesized that if these signatures are indeed artifactual, then their removal from MDA neurons would result in sSNV accumulation rates more consistent with PTA neurons. Indeed, after subtracting the Signature B-like exposure from MDA neurons, the yearly accumulation rate by SCAN2 decreased from 25.7 sSNVs/year to 16.7 sSNVs/year, more closely matching that of PTA neurons (**Supplementary Fig. 11**). Taken together, these observations provide compelling evidence that sSNVs accumulate in human neurons at a rate closer to 15 sSNVs/year and that Signature B consists largely of MDA technical artifacts.

Finally, we emphasize that although a majority of SCAN2’s calls in infant PTA neurons are C>Ts, they are materially different from those found by SCAN2 in MDA neurons and are more likely to be true mutations. This is easily seen upon computing enrichment for C>Ts by normalizing by the frequencies of N<u>**C**</u>N trinucleotide contexts in the human genome (Methods). After normalization, PTA C>Ts show a clear and strong preference for <u>**C**</u>pG contexts in a manner similar to COSMIC signature SBS1 (**Fig. 2f-i**, right panel), a mitotic clock-like signature believed to occur during cell division^28^. This suggests cell division during embryogenesis and subsequent development as plausible sources for infant PTA C>Ts. Among the normalized MDA spectra, a similar but smaller bias toward <u>**C**</u>pG contexts exists in the infant MDA calls and Signature B but not in Signature scF. These data suggest that neurons in the infant brain contain lower levels of single-neuron sSNVs than previously reported, but, since we remove any sSNV present in matched bulk, also underestimates the number of clonal sSNVs in neurons which are likely to number in the hundreds^29^.

### Characteristics of somatic indels in single human neurons

SCAN2 provides the first catalog of somatic indels from single cells and the first such catalog from a post-mitotic human cell. In total, 532 indels were identified from the 25 PTA-amplified neuronal genomes. Somatic indels increased with age by 2 to 4 somatic indels per neuron per year (Methods, **Fig. 3a**), which is surprisingly similar to rates observed in several mitotically active cell types^8–10,30^. However, we caution that these rates are difficult to calculate for the reasons explained above: indel sensitivity is highly dependent on indel length and genomic context and, in particular, our method has low sensitivity for highly mutable sites such as microsatellites that may recur in multiple individuals. We therefore propose a rate of ^~^2 somatic indels per year as a lower bound. Deletions accumulated 3.3-fold faster than insertions (**Fig. 3b**) and indel sizes ranged from −28 bp to +14 bp (**Fig. 3c**). As was the case for sSNVs, MDA yields a higher accumulation rate of 3.0 somatic indels/year and we again attribute this increase to MDA artifacts; MDA somatic indels are not included in the following analyses.

**Figure 3:**
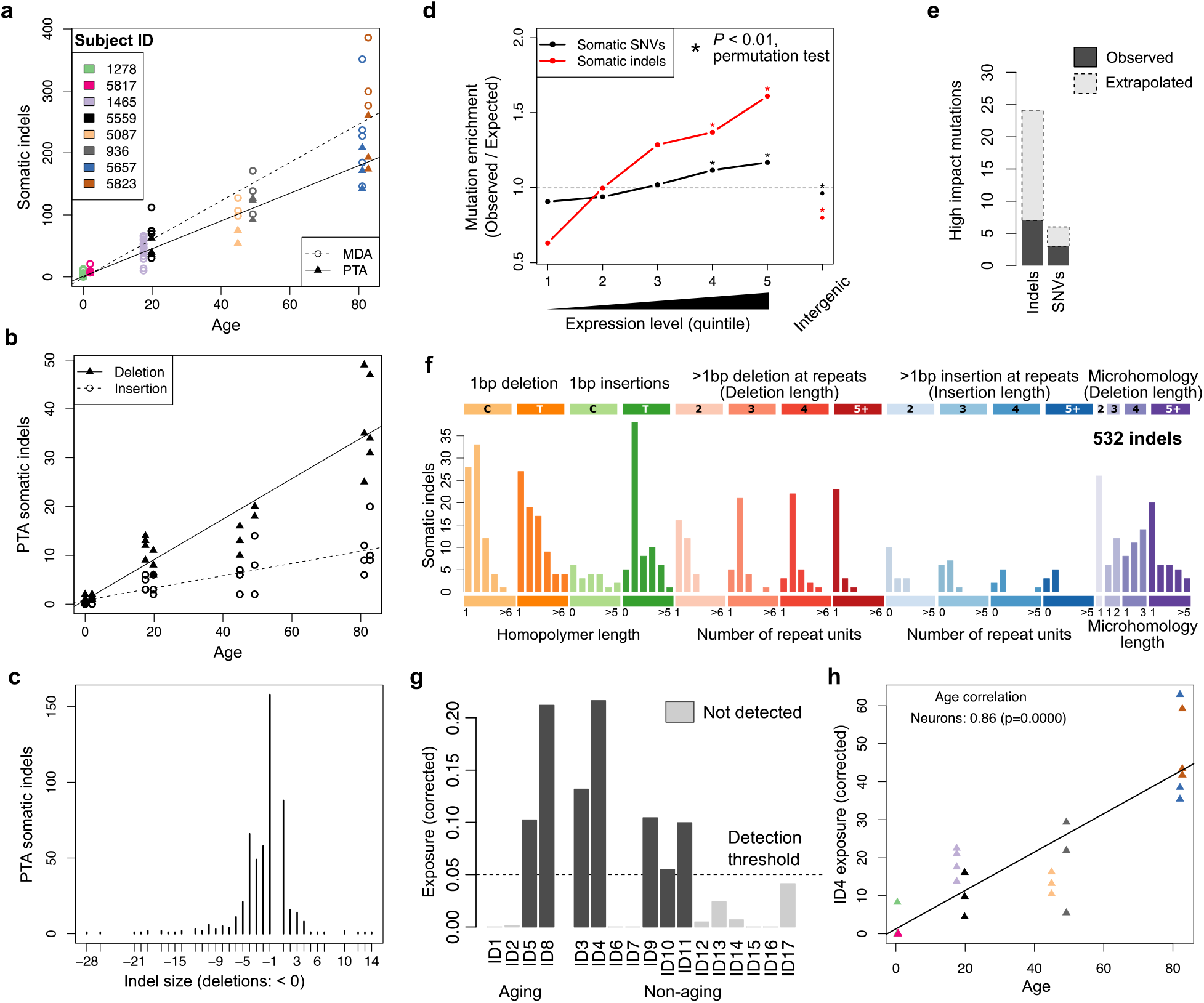
Characteristics of somatic indels in aging human neurons. **a.** Age-related increase of somatic indel burden across 8 individuals. Adjustment for sensitivity shown here represents the lower bound corresponding to ~2 somatic indels per neuron per year. **b.** Age-related increase of somatic insertions and deletions called from PTA neurons, separately. **c.** Distribution of somatic indel lengths from PTA neurons. **d.** Enrichment of PTA somatic SNVs in intergenic regions and transcribed regions stratified by expression quartile. Expression levels were derived from GTEx and range from quintile 1 (lowest) to quintile 5 (highest). Expected number of mutations determined by permutation testing (*: p < 0.01). **e.** Number of high impact mutations according to SnpEff (dark grey); expected number of high impact mutations after adjusting for sensitivity (light grey). **f.** Mutation spectrum identified by de novo signature extraction from 532 somatic indels. **g.** Exposures to COSMIC ID signatures calculated by least squares fitting. Exposures were corrected by normalizing indel counts by ID83 channel-specific sensitivity (**Supplementary Fig. 8c**) before fitting. **h.** Age association of ID4, a signature of unknown aetiology, with neuron age.

Similar to sSNVs, somatic indels occur more frequently in genic regions, and the enrichment for both forms of mutation is significantly increased in highly transcribed genes (**Fig. 3d**). Of the 22 exonic indels detected, 7 were scored as high impact (frame shift mutations in TIA1, MYO3B, PASK, CCDC162P, ZSCAN32, FAM161B, and CHSY1); in contrast, only 3 sSNVs were scored as high impact (stop gain in ZDHHC12, structural interaction change in PIP4K2B and a splice acceptor mutation in ANGPTL4). After adjusting for detection sensitivity, 24 high severity somatic indels and 6 high severity somatic SNVs would be expected to exist in the PTA cohort (**Fig. 3e**), suggesting that indels may have an equal or greater functional impact compared to sSNVs despite accumulating at an ^~^8-fold lower rate.

*De novo* mutation signature extraction yielded only a single ID83 somatic indel spectrum, likely due to the limited number of somatic indels (**Fig. 3f**), that resembles spectra from dividing cells^9,10,30^ (**Supplementary Fig. 12**). After correcting for ID83 class-specific sensitivity, fitting to the COSMIC signature catalogue and removing signatures with <5% contribution, 7 indel signatures were detected, including two clock-like signatures ID5 and ID8 (**Fig. 3g, Supplementary Fig. 13**). The two remaining clock-like signatures ID1 and ID2 were not detected, consistent with the facts that neurons are post-mitotic and that the proposed aetiology for ID1 and ID2 involves DNA replication. The most prevalent signature was ID4: a signature observed in several cancer types but with no proposed mechanism. Surprisingly, ID4 is more strongly correlated with age in neurons than the clock-like signatures ID5 and ID8 (**Fig. 3h**; correlation with age = 0.86, 0.53 and 0.72, respectively). ID3 was recently detected in normal bronchial epithelium^30^, especially in smokers, and also shows correlation with age in neurons (correlation = 0.73). The remainder of the detected signatures (ID9, ID10 and ID11) are relatively poorly correlated with age and may represent artifacts of the signature fitting process.

## Discussion

It is now clear that MDA genome amplification can suffer from single-stranded dropout, creating C>T artifacts that are often indistinguishable from mutations. These artifacts can be separated out by mutation signature analysis in some applications: for example, we successfully identified an sSNV signature that increases with age in human neurons despite the presence of these MDA artifacts^6^ and confirmed this signature using PTA. Further, the similarity between SNV accumulation rates from PTA cells and MDA cells after subtracting signature B suggests that an improved correction method may be able to accurately estimate total mutation burdens from MDA. PTA introduces fewer artifacts due to its quasilinear amplification process and offers the ability to call individual mutations with high specificity. However, even using PTA, cells with low mutation burdens must be analyzed by highly specific genotypers aware of single-cell amplification artifacts.

The methods introduced in SCAN2 come with important caveats. First, the multi-sample sSNV calling approach must be applied to batches of PTA-amplified single cells that have been exposed to similar mutational processes. Further, the efficacy of the multi-sample mutation signature approach depends on the similarity between the true signature under study and the universal PTA artifact signature: higher similarity will yield fewer benefits. The worst-case scenario occurs when the two signatures are identical; under these circumstances multi-sample calling would yield no improvement. Somatic indel detection depends on a sufficiently large sample set for screening recurrent artifacts. Notably, this filtration strategy is expected to limit SCAN2’s ability to detect somatic indels at highly mutable sites such as microsatellites.

In this study we examine indels in post-mitotic single cells for the first time. Because these cells no longer divide, the active mutational processes must not be associated with DNA replication. This may help to narrow down the possible mechanisms underlying indel signatures ID4 and ID5, whose aetiologies remain unknown. Transcriptionally associated mechanisms are the clearest candidate for further inquiry due to the enrichment of indels in expressed genes^31^; however, larger datasets are needed to draw conclusions with confidence.

## Methods

### Human tissue and case selection

Postmortem frozen human tissues were obtained from the NIH Neurobiobank at the University of Maryland School of Medicine. Samples were obtained and processed according to IRB-approved protocol. Non-disease neurotypical individuals had no clinical history of neurologic disease and were selected to represent a range of ages from infancy to older adulthood.

### Isolation of single neuronal nuclei for single-cell whole genome sequencing

Single neuronal nuclei were isolated using fluorescence-activated nuclear sorting (FANS) for NeuN, as described previously^6,32^. Briefly, nuclei were prepared from unfixed frozen human brain tissue, previously stored at −80°C, in a dounce homogenizer using a chilled tissue lysis buffer (10mM Tris-HCl, 0.32M sucrose, 3mM Mg(OAc)2, 5mM CaCl2, 0.1mM EDTA, 1mM DTT, 0.1% Triton X-100, pH 8) on ice. Tissue lysates were carefully layered on top of a sucrose cushion buffer (1.8M sucrose 3mM Mg(OAc)2, 10mM Tris-HCl, 1mM DTT, pH 8) and ultra-centrifuged for 1 hour at 30,000 × g. Nuclear pellets were incubated and resuspended in ice-cold PBS supplemented with 3mM MgCl2, filtered (40 μm), then stained with Alexa Fluor 488-conjugated anti-NeuN antibody (Millipore MAB377X). Large neuronal nuclei were then subjected to FANS, one nucleus per well into 96-well plates.

### Single nucleus whole genome amplification by primary template-directed amplification (PTA)

Isolated single neuronal nuclei were lysed and their genomes amplified using PTA, a recently developed method that pairs an isothermal DNA polymerase with a termination base^15^. PTA reactions were performed using the ResolveDNA EA Whole Genome Amplification Kit (formerly SkrybAmp EA WGA kit) (BioSkryb, Durham, NC), using the manufacturer’s protocol. Briefly, single nuclei were sorted into wells containing 3 μL Cell Buffer pre-chilled on ice, then alkaline lysed on ice with MS Mix, mixed at 1400rpm, then neutralized with SN1 Buffer. SDX buffer was then added to the neutralized nuclei followed by a brief incubation at room temperature. Reaction-Enzyme Mix were added, then the amplification reaction was carried out for 10 hrs. at 30°C, followed by enzyme inactivation at 65°C for 3 min. Amplified DNA was then cleaned up using AMPure, and yield determined by the picogreen method (Quant-iT dsDNA Assay Kit, ThermoFisher). Samples were subjected to quality control by multiplex PCR for 4 random genomic loci as previously described^6^, and by Bioanalyzer for fragment size distribution. Amplified genomes demonstrating positive amplification for all 4 loci were then prepared for Illumina sequencing.

### Library preparation for scWGS

Libraries were made following a modified KAPA HyperPlus Library Preparation protocol provided in the ResolveDNA EA Whole Genome Amplification protocol. Briefly, end repair and A-tailing were performed for 500 ng of amplified DNA. Adapter ligation was then performed using the SeqCap Adapter Kit (Roche, 07141548001). Ligated DNA was cleaned up using AMPure and amplified through an on-bead PCR amplification. Amplified libraries were selected for 300-600 bp size using AMPure. Libraries were subjected to quality control using picogreen and Tapestation HS D1000 Screen Tape (Agilent PN 5067-5584) before sequencing. Single cell genome libraries were sequenced on the Illumina NovaSeq platform (150bp × 2) at 30× except for subjects 1278 (HiSeq, 60×) and 1465 (NovaSeq, 60×).

### Single-cell amplification quality metrics

Median absolute pairwise differences (MAPD) were computed by estimating copy number in bins *CN*_*i*_ of size 50 kb following ref. 33; subsequently, MAPD = median(|log_2_ *CN*_*i*_ − log_2_ *CN*_*i*+l_|). Copy number profiles in **Fig. 1** were produced using Ginkgo^34^ with bin size 100 kb, variable binning enabled and pseudoautosomal regions masked. Allele balance distributions were computed separately for each cell by measuring single-cell VAFs at all heterozygous SNP sites used to train the SCAN2 allele balance model and then applying R’s `density` function.

### Large somatic copy number alteration analysis

Large-scale somatic CNA analysis used Ginkgo with variable bin size=1 Mb to produce a profile of normalized read counts for all bulks in PTA single cells. Large somatic CNA candidates were defined as runs of 5 or more windows *i* with read depth ratio *S_j,i_*/*B_i_* < 0.6 or > 1.4, where S*_j,i_* denotes the normalized read depth in window *i* in single cell *j* and *B_i_* is the same normalized window in the matched bulk sample. Further, somatic CNA candidates were required to have neutral copy number in the matched bulk by the same metrics. This CNA calling procedure is crude and only intended to recover very large (>5 MB) CNAs; however, these parameters successfully recovered male X chromosomes and female Y chromosomes in bulk and the large deletions observed in the PTA-amplified neuron 5823PFC-B (**Supplementary Figure 2**). Apart from 5823PFC-B, no autosomal somatic CNAs were detected by this method.

### Somatic mutation calling on male X chromosomes

GATK HaplotypeCaller (v3.8.1) was run in joint mode across all samples (bulk, PTA and MDA) for each individual using dbSNP 147_b37_common_all_20160601 and parameters `--dontUseSoftClippedBases -rf BadCigar -mmq60`. Pseudoautosomal regions were not included. The resulting VCF was filtered for SNVs using GATK SelectVariants `- selectType SNP-selectType INDEL -restrictAllelesTo BIALLELIC - env -trimAlternates`. Somatic SNVs and indels in single cells were called separately using the following criteria: VAF > 90%, single cell depth > median(single cell depth), 0 alternate reads in bulk, bulk depth > 10 and absence from dbSNP. A set of germline SNPs and indels for estimating sensitivity was defined by sites with bulk VAF > 90%, bulk depth > median(bulk depth) and no more than 2 reference reads in bulk. For each single cell, the fraction of these sites passing the somatic filters (except for requiring 0 alternate reads in bulk and absence from dbSNP) was used as an estimate of somatic mutation sensitivity. The final estimated number of mutations was calculated by (corrected calls) = (#somatic mutations called) / (estimated sensitivity). Excess MDA calls were called per individual as the median(corrected MDA calls) – median(corrected PTA calls).

### sSNV false discovery rate estimation

Estimated FDR curves shown in **Figure 2a** were parameterized by

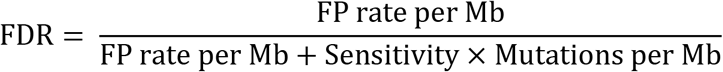

Parameters used were: PTA with GATK (ref. 15), FP rate per Mb = 0.9, sensitivity = 0.8; PTA with SCAN2 (multi-sample calling) FP rate per Mb = 0.0143, sensitivity = 0.457 (derived from simulation experiments, see Synthetic diploid simulations). To compute the best-case scenario for MDA, we assumed that all artifacts caused by single stranded dropout would be erroneously identified as true SNVs and that these would be the only source of FPs. The number of single-stranded dropout artifacts in MDA was estimated by the excess number of sSNV calls per hemizygous X chromosome (15.9 sSNVs). To convert to FPs per diploid megabase, the excess rate is first doubled and then divided by 152,231,524 bp, the size of chromosome X after removing pseudoautosomal regions. This yielded a rate of 0.21 FPs per Mb, which was applied to the whole genome. Finally, because these FPs should be called with similar sensitivity to true mutations, there was no need to provide a sensitivity parameter for the best-case MDA scenario since it would cancel out in the above equation.

### Multi-sample somatic SNV calling procedure with SCAN2

First, a set of high quality somatic SNV calls is produced for each single cell by running SCAN-SNV in single sample mode (as described in ref. 14) with a stringent target FDR of 1%. The true sSNV mutation spectrum is then produced by combining calls from all 25 PTA cells into a single, raw SBS96 mutation spectrum. In general, this multi-sample combination step should only be applied to cells exposed to the same mutational process (e.g., treatment by the same chemical mutagen). Exposures to the true spectrum and universal PTA artifact spectrum (described below) are computed for each single cell by least squares fitting. Weights are computed for each cell *i* and rejected sSNV candidate *j* using a likelihood ratio

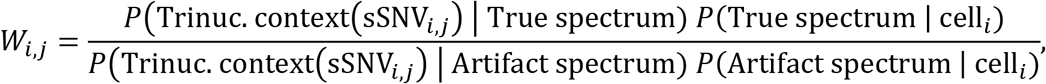

where *P*(Trinucleotide context(sSNV_*j*_)| True spectrum) is the component of the true mutation spectrum corresponding to the mutation type and context of sSNV*_j_* and *P*(True spectrum | cell_*i*_) is cell *i*’s estimated exposure to the true mutation signature. The same meanings apply to the artifact spectrum. Therefore, *W_i,j_* > 1 indicates lower likelihood of sSNV*_i,j_* being produced by the artifact process while *W_i,j_* < 1 indicates higher likelihood. The weight is used to adjust a previously described heuristic^14^ that estimates the ratio of true mutations *N_T_* and artifacts *N_A_* among candidate sSNVs with similar VAF and sequencing depth as the candidate sSNV being evaluated. This produces a multi-sample adjusted, Phred-scaled quality score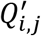:

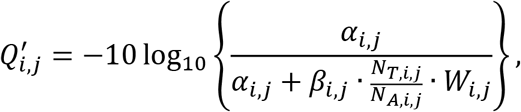

where *α*_*i,j*_ and *β*_*i,j*_ are the type I error rate and power for sSNV*_i,j_* estimated by the pre-amplification artifact model used by SCAN-SNV (ref. 14 provides more details on this model). Finally, the rejected candidate sSNV*_i,j_* is accepted if it was previously rejected only by the pre-amplification artifact model (i.e., passing all other criteria from ref. 14) and 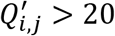, corresponding to a desired FDR of 1%. This threshold can be set by the user.

### Estimation of genome-wide somatic SNV burden

In addition to providing a set of sSNV calls, SCAN2 also estimates the genome-wide somatic SNV burden by estimating sSNV detection sensitivity at a subset of the high confidence, heterozygous germline SNPs (hSNPs) used to train the allele balance model. First, SCAN2 calculates the distance to the nearest training hSNP for all candidate somatic SNVs and forms the distribution of these distances. The training set of germline hSNPs is then downsampled, using importance sampling, so that the distribution of distances to the nearest hSNP matches that of somatic SNV candidates. This step is necessary because the accuracy of the spatial allele balance model increases as distance to the nearest hSNP decreases. Once the downsampled set of germline hSNPs is selected, each hSNP is individually analyzed using a leave-1-out approach: the hSNP is removed from the allele balance training set, the model predicts the allele balance at the hSNP and the hSNP is then assessed using all somatic calling criteria except for dbSNP exclusion and lack of supporting reads in bulk. Only hSNPs that meet the depth requirements for somatic calling (set by the user; default: sequencing depth of the matched bulk > 10 and depth in the single cell > 5) are assessed. Among these, the fraction *f_h_* of hSNPs passed by the somatic caller serves as an estimate of somatic sensitivity. The rate of somatic SNVs per haploid gigabase is then

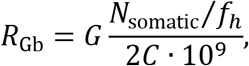

where *C* is the number of diploid gigabases of the genome with sufficient sequencing depth for analysis, as specified by the user, and is collected by GATK DepthOfCoverage at base pair resolution. *G* is the total genome size; for **Figure 2d**, *G*=5.845 corresponds to the number of autosomal haploid gigabases and matches ref. 6; for synthetic diploid simulations, *G*=0.3044, corresponding to twice the size of the haploid, non-pseudoautosomal region of chromosome X in GRCh37. **Supplementary Figure 6d** provides an assessment of the accuracy of this estimate in simulated data with known mutation burdens.

### Deriving the universal PTA artifact spectrum

The universal PTA artifact spectrum was derived in 2 steps (technical details are provided in the next paragraph). First, two sets of sSNVs enriched for artifacts were extracted for each male sample (**Supplementary Fig. 3a)**: (1) *S*_X artifact_ from X chromosomes (male samples only) and (2) *S*_Autosomal artifact_ from autosomal SNV candidates with VAFs consistent with expectation for pre-amplification artifacts, as determined by the local allele balance. *S*_Autosomal artifact_ was added because *S*_X artifact_ consisted of only 190 likely artifacts, which may be insufficient to produce a high quality 96-dimensional mutation spectrum. Second, *de novo* signature extraction was performed on *S*_X artifact_, *S*_Autosomal artifact_ and an additional set *S*_PASS_ of high quality sSNVs (**Supplementary Fig. 3b)**. The high quality sSNV set provides the true mutational signature, helping to prevent true mutations in *S*_X artifact_ or *S*_Autosomal artifact_ from being assigned to the artifact signature. De novo signature extraction produced N=2 signatures, as expected: one corresponding to *S*_PASS_ and a second corresponding to the PTA high-VAF artifact process, which became the universal PTA artifact spectrum (**Supplementary Fig. 3c**). Estimated exposures to the true and artifact spectra confirmed that the two artifact sets were highly enriched for artifacts, contrasting with the high-quality set (**Supplementary Fig. 3d**). The similarity between the PTA universal artifact signature and the MDA artifact C>T signature is notable and provides evidence that the signature is unlikely to be an overfit to this dataset.

In more detail, X chromosome artifacts were identified from candidate SNVs produced by GATK HaplotypeCaller (as described in *Somatic mutation calling on male X chromosomes*) by requiring the SNV candidate to: (1) occur in the non-pseudoautosomal X regions, (2) have total sequencing depth >= median(sequencing depth) of the X chromosome, (3) be supported by at least 6 alternate reads, and (4) have 35% <= VAF <= 75%. Autosomal artifacts were identified by the SCAN2 allele balance consistency (ABC, *P*_true_) and pre-amplification test (*P*_artifact_) *P*-values (see ref. 14). Briefly, large ABC *P*-values indicate that the candidate SNV’s VAF is consistent with the locally estimated allele balance, as should be the case for a true mutation. Large pre-amplification *P*-values indicate that the candidate’s VAF is consistent with that expected for an early-occuring artifact. Autosomal SNV candidates which fail the pre-amplification test, pass all other SCAN2 tests and for which *P*_amplification artifact_ > *P*_ABC_ were selected as autosomal artifacts. *S*_PASS_ is the set of SNVs called by SCAN2 in single sample mode using the stringent calling parameter `--target.fdr=`0.01 (i.e., PASS sSNVs). *De novo* signature extraction was performed by SigProfiler^35^ version 2.5.1.7, as used in other *de novo* extractions. Signature channels with values < 10^−4^ were replaced by 10^−5^ to prevent channels with extreme weights.

### Somatic indel detection with SCAN2

Candidate somatic indels are initially constructed by GATK HaplotypeCaller using the same parameters as in section *Somatic mutation calling on male X chromosomes.* Somatic indels are assessed by all tests and filters applied to somatic SNVs in standard single-sample mode and an additional single-cell depth requirement of 10 reads. Notably, the allele balance model applied to candidate somatic indels is not built using germline indels; rather, the same model trained on germline hSNPs and applied to sSNVs is used for indel calling. Somatic indels passed by this process are then filtered using the cross-sample site list by requiring either: (1) reads supporting the somatic indel exist only in single cells from one individual or (2) no single cell contains more than 2 supporting reads, regardless of the number of cells and subjects in which these indel-supporting reads appear. The cross-sample list is generated by running GATK HaplotypeCaller (with the same parameters as in indel discovery) jointly on whole-genome amplified single cells from at least two individuals. Multi-sample mutation signature calling is not applied to indels, although it may be found to be beneficial with further development.

### Synthetic diploid simulations

Synthetic X diploids (SDs), as described in ref. 14, were used to assess the performance of SCAN2. Briefly, synthetic X diploids are constructed by merging chromosome X-mapped sequencing reads from two male, independently amplified single cells. This process creates a reasonably accurate amount of allelic amplification balance and amplification artifacts. In this study, 9 SDs with 30× mean depth were generated by making all pairings of the 3 PTA cells from donor 1278 and 3 PTA cells from donor 5817. The youngest donors (0.4 and 0.6 years old) were chosen to minimize the number of true somatic mutations endogenous to each X chromosome prior to adding spike-in mutations. To identify somatic SNVs endogenous to each X chromosome, GATK HaplotypeCaller was applied jointly to the SDs and the 7 PTA donor cells using the same parameters as in *Somatic mutation calling on male X chromosomes*. An additional HaplotypeCaller run using `-mmq 1` was also performed. Endogenous sSNVs were identified by applying the following hard filters: VAF=100% or VAF >= 90% with fewer than 2 reference reads; depth >= 5 in the single cell, depth > 10 in the matched bulk and no mutation supporting reads in bulk in either the mapping quality 60 or mapping quality 1 runs. A single cluster of sSNVs at chrX:77471371-77471423 that appeared to be caused by clipped alignment was manually removed from the endogenous somatic mutation list. No endogenous indels were identified.

Each SD received 20, 50, 100, 200, 500, 1000 and 2000 spike-ins, evenly split between SNVs and indels, for a total of 63 SDs. SDs with 1000 and 2000 spike-ins were used only for the rate estimation analysis presented in **Supplementary Figure 6d**. Somatic SNV spikeins were randomly generated as previously described^14^. Somatic indel spikein candidates were randomly generated until ^~^1000 candidates were obtained for each ID83 class. Indel ID83 classes were determined by first left-aligning indels by `bcftools norm` and then using SigProfilerMatrixGenerator^36^ to assign ID83 status. Somatic indel spikeins were required to be at least 150 bp away from the nearest indel spikein candidate to prevent crowding in repetitive tracts and potential alignment issues caused by clustered indels. SNV and indel spikeins were not allowed to overlap. SCAN2 was run jointly on the set of 63 SDs with the same parameters used in the analysis of single neurons. Sensitivity was calculated by the fraction of successful spike-ins recovered; any SNV call not in the endogenous sSNV or spike-in sets was considered a false positive. Due to the ambiguous nature of indel representation, indel calls were considered matches to known spike-ins if either: (1) the calls matched the spike-in indel exactly or (2) the called indel was the correct length and was located exactly 1 bp away from the spike-in location.

### SNV calling with Monovar

Monovar commit `7b47571` was downloaded and the somatic calling strategy reported previously^22^ was mimicked as closely as possible, using scripts developed in ref. 14 (N.B., the authors provide no script for identifying somatic mutations). Single cell BAMs were input to samtools version 1.9 with options `-BQ0 -d10000 -q 40`, which was piped into the monovar.py script with options `-p 0.002 -a 0.2 -t 0.05 -m` 2 as recommended by the authors. To determine whether SNVs were somatic or germline, samtools was run with the same options on matched bulk data. Somatic SNVs were determined by the following filters: Monovar’s genotype string must not match `./. or 0/0;` a minimum sequencing depth of 10 in the single cell with at least 3 reads supporting the mutation; at least 6 reads in bulk with no more than 1 mutation supporting read; and single cell VAF ≥ 10% for sSNVs with >100 depth or VAF ≥ 10% for sSNVs with depth between 20 and 100. Finally, sSNVs were filtered if any other call occurred within 10 bp.

### SNV calling with SCcaller

SCcaller version 1.1 was run as previously reported^26^, using scripts developed in ref. 14. BAMs were converted to pileups using samtools version 1.3.1 with the option -C50 and hSNPs were defined using dbSNP version 147 common. Single cell somatic SNVs were called by applying SCcaller’s `-a varcall, -a cutoff` and reasoning v1.0 script in sequence with default parameters. As recommended on SCcaller’s Github README, passing somatic mutations were required to have VAF > 1/8, filter status `= PASS`, bulk status `= refgenotype` and must not have been observed in dbSNP. The standard calling parameter is *α* = 0.05, while the stringent calling parameter is *α* = 0.01.

### SNV calling with LiRA

LiRA version `1f4cab4` was run following instructions on Github. The joint VCF produced internally by SCAN2 (/path/to/scansnv/gatk/hc_raw.mmq60.vcf) for each individual was supplied as the input VCF to LiRA. All samples were processed as male regardless of sex to restrict calls to the autosomes and to use a single consistent genome size for total burden estimation (LiRA accounts for the difference in genome size between males and females due to chrY). LiRA uses a genome size *G*=6.349 for males (see *Estimation of genome-wide somatic SNV burden*); to restrict to autosomal extrapolation (*G*=5.845) as used in all other sections and in ref. 6, LiRA total SNV burden estimates were multiplied by 5.845/6.349. LiRA total burden estimates retrieved from ref. 6, supplementary table S5 were not corrected in this way since they were already computed using *G*=5.845.

### Somatic SNV analysis of single human neurons

MDA and PTA single neurons were analyzed by SCAN2 with identical parameters. Non-default parameters: `--abmodel-chunks=4, --abmodel-samples-per-chunk=5000, --target-fdr=0.01;` data resources: human reference genome GRCh37d5, SHAPEIT phasing panel 1000GP_Phase3 and dbSNP version 147_b37_common_all_20160601. SCAN2 was run jointly on MDA and PTA cells for each subject, but subjects were analyzed in separate runs (8 total SCAN2 runs corresponding to 8 subjects). Notably, even single-sample SCAN2 uses joint GATK HaplotypeCaller to create the initial set of candidate somatic SNVs, though additional information shared across cells is not used in single-sample mode. Multi-sample SCAN2 was run jointly on the SCAN2 results for all 25 PTA samples from the per-subject SCAN2 runs. sSNV accumulation rates with age were derived from a mixed-effects linear model that accounts for the fact that multiple neurons from the same individual are not independent measurements, as would be assumed by a simple linear regression. Mixed-effects model fitting was performed using the `lme4 R package with the command lmer(age ~ total_burden + (1|subject))`, where `total_burden` refers to the genome-wide burden estimate described in *Estimation of genome-wide somatic SNV burden*.

Mutation spectra in **Figures 2f,g** are the counts of passing sSNVs from high-confidence, single-sample SCAN2 over samples 1278BA9-A, 1278BA9-B, 1278BA9-C, 5817PFC-A, 5817PFC-B and 5817PFC-C (infant PTA) and samples 1278_ct_p1E3, 1278_ct_p1E6, 1278_ct_p1G9, 1278_ct_p2B9, 1278_ct_p2C7, 1278_ct_p2E4, 1278_ct_p2E6, 1278_ct_p2F5, 1278_ct_p2G5, 5817_ct_p1H10, 5817_ct_p1H2, 5817_ct_p1H5 and 5817_ct_p2H6 (infant MDA). Multi-sample mode should not be used for mutation signature analysis since it is biased against SBS96 channels that contribute to the universal artifact signature. To normalize for hg19’s trinucleotide content, all 3mers (including overlaps) were extracted from the primary autosomal contigs in GRCh37 and tabulated. Each SBS96 channel was divided by the frequency of the associated 3mer in hg19.

### Removal of signature B from MDA samples

Signature B levels in MDA samples were measured by de novo signature extraction from the combined set of 76 PTA and MDA neurons using `SigProfiler` version 2.5.1.7. 3 signatures were discovered, with one nearly identical to signature B^6^ (cosine similarity=0.996). Removal of signature B as shown in **Supplementary Figure 11** was achieved by subtracting the reported number of sSNVs attributed to signature B from the total number of called sSNVs in each MDA sample.

### Somatic indel analysis of single human neurons

SCAN2 was run on PTA with the same parameters used in SNV analysis (most notably, `--target.fdr=0.01)`. The cross-sample filtration list was generated using all 76 MDA and PTA single cells analyzed in this study. Indels were classified into ID83 channels using `SigProfilerMatrixGenerator`. For MDA-amplified neurons only, somatic indels were additionally filtered to remove all single base insertions in homopolymers of length 3 or greater (i.e., ID83 classes 1:Ins:C: 3-5 and 1:Ins:T:3-5). Somatic indel sensitivity was computed in two ways following the process in *Estimation of genome-wide somatic SNV burden*. First, germline heterozygous indels discovered in bulk were downsampled to match the ID83 spectrum of called somatic indels to provide a set of indels with roughly similar characteristics to somatic indels. Total sensitivity was computed on the downsampled germline set, giving a sensitivity-adjusted *N*_somatic_ = (# called indels) / (germline sensitivity). Since the cross-subject panel was not applied to the germline heterozygous indels (because they are common polymorphisms and are often shared), this overestimates sensitivity and underestimates of the number of indels. Second, the ID83 channel-specific sensitivities derived from SD simulations were applied to each single cell individually by dividing the number of somatic indels calls per channel by the channel-specific sensitivity. Summing over all ID83 classes gives *N*_somatic_ per cell. The final rate *R*_Gb_ was estimated as explained above. De novo extraction was performed by `SigProfiler` on PTA neurons only, which produced only a single signature. Fits to COSMIC indel signatures were performed using the COSMIC version 3 set of indel signatures ID1-18. For the discovery of active signatures in **Figure 3g**, all 532 indels were combined into a single set and exposures to each of the 17 signatures were estimated by least squares fitting using `lsqnonneg` from the `pracma` R package. Otherwise (e.g., for analysis of correlation with age), somatic indels were kept separate and fit using the same method.

### Functional impact of point mutations

The severity of somatic SNV and indel mutations reported in Figure 3 were derived from SnpEff version 4.3t using the hg19 database. High and moderate mutations were those annotated as HIGH or MODERATE, respectively, in the first reported annotation field. The genes impacted and protein-altering effects were also taken from the first annotation field. The extrapolation from called mutations to the expected number over the PTA cohort used cohort-wide sensitivity estimates of 47.7% for sSNVs and 29.3% for somatic indels corresponding to (# SCAN2 PTA sSNV calls = 7174) / (sum of estimated PTA sSNV burdens = 15,030) and (# SCAN2 PTA indel calls = 532) / (germline sensitivity-based estimate for total indel burden = 1812), respectively. The number of expected high-impact mutations per cohort is the number of observed HIGH impact mutations (n=7, indels, n=3, sSNVs) divided by sensitivity.

### Enrichment of somatic mutations in transcribed genes

Gene expression quantification data were obtained from the GTEx consortium (version 8); gene start and stop genomic positions were obtained by matching GENCODE v26 (hg38 to b37 liftover) “gene” records (column 3) to the GTEx expression matrix using Ensembl gene IDs. Autosomal genes with mean TPM > 1 in either 209 frontal cortex (BA9) GTEx samples or across the full GTEx dataset were retained for analysis. Genes retained by this filtration were then ranked by average TPM across the 209 BA9 samples and separated into expression quintiles. Each somatic mutation was assigned to 1 of 6 bins (5 expression quintiles and intergenic) based on overlap with this gene set. Mutations overlapping multiple genes were resolved by assigning the mutation to the first gene in the overlap list. Enrichment analysis was performed by permutation: for each single cell, mutation positions were randomly shuffled across the genome 250 times to create a null distribution of mutation density. To approximate calling sensitivity, position shuffling was restricted to the subset of each single cell genome that met the minimum depth requirements for SCAN2 analysis. To perform enrichment calculations, observed mutation counts for each bin *i* were combined across all samples for either the observed data *D_i_* (7,174 sSNVs by multi-sample SCAN2 or 532 indels, separately) or one of the shufflings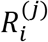, *j* = 1 to 250. Enrichment levels were calculated for observations and permutations by dividing each bin count by the mean count over the 250 shufflings 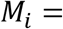 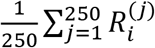. Two-tailed *p*-values were determined for each bin *i* by counting the fraction of permutations with absolute log ratios exceeding the observed absolute log ratio:

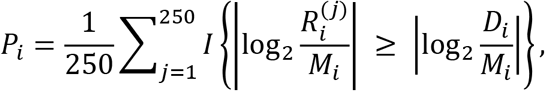

where *I*(·) is the indicator function.

## Data availability

All MDA-amplified single neurons and matched bulks listed in **Supplementary Table 2** were downloaded from dbGaP, identifier phs001485.v1.p1. Only neurons from the pre-frontal corteces from individuals for which additional PTA data were generated were used. *Raw sequencing read data for PTA-amplified single cells will be uploaded to dbGaP.*

## Code availability

SCAN2 is available for download at https://github.com/parklab/SCAN2.

---

**Supplementary Figure 1:**
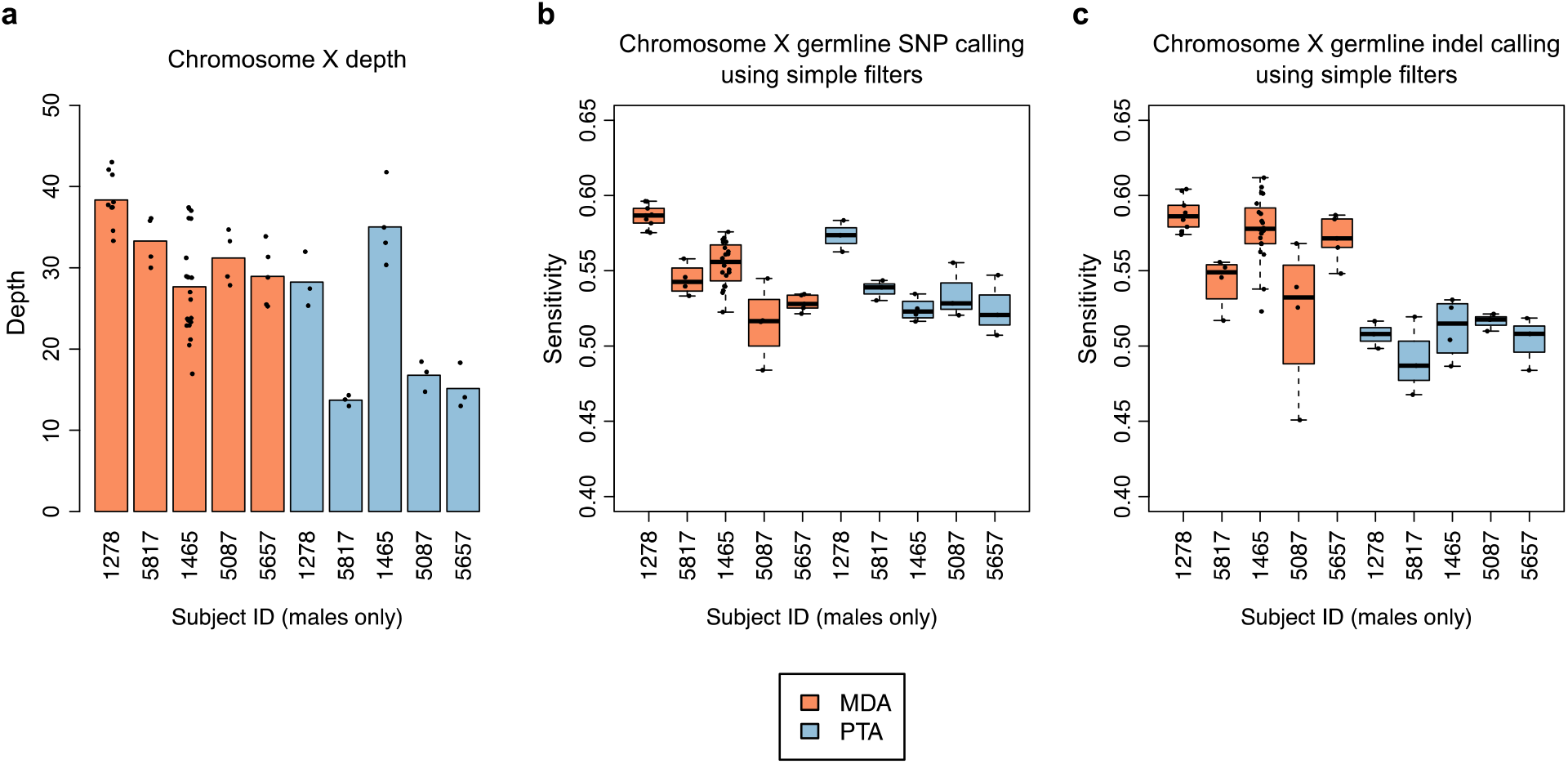
Simple somatic mutation calling on male chromosome X. **a.** Mean sequencing depth per cell (points) and averaged over all cells per donor (bar). PTA cells for subjects 1278 and 1465 were sequenced to ~60× total depth while other PTA cells were sequenced to ~30×. Chromosome X in males should be sequenced to about half of the genome-wide mean depth due to hemizygosity. **b.** Sensitivity for germline SNPs using somatic SNV calling criteria (depth and allele fraction filters). Germline SNP sensitivity provides an estimate for somatic SNV sensitivity. **c.** Same as (b) for indels. Boxplot whiskers, furthest point at most 1.5× interquartile range.

**Supplementary Figure 2:**
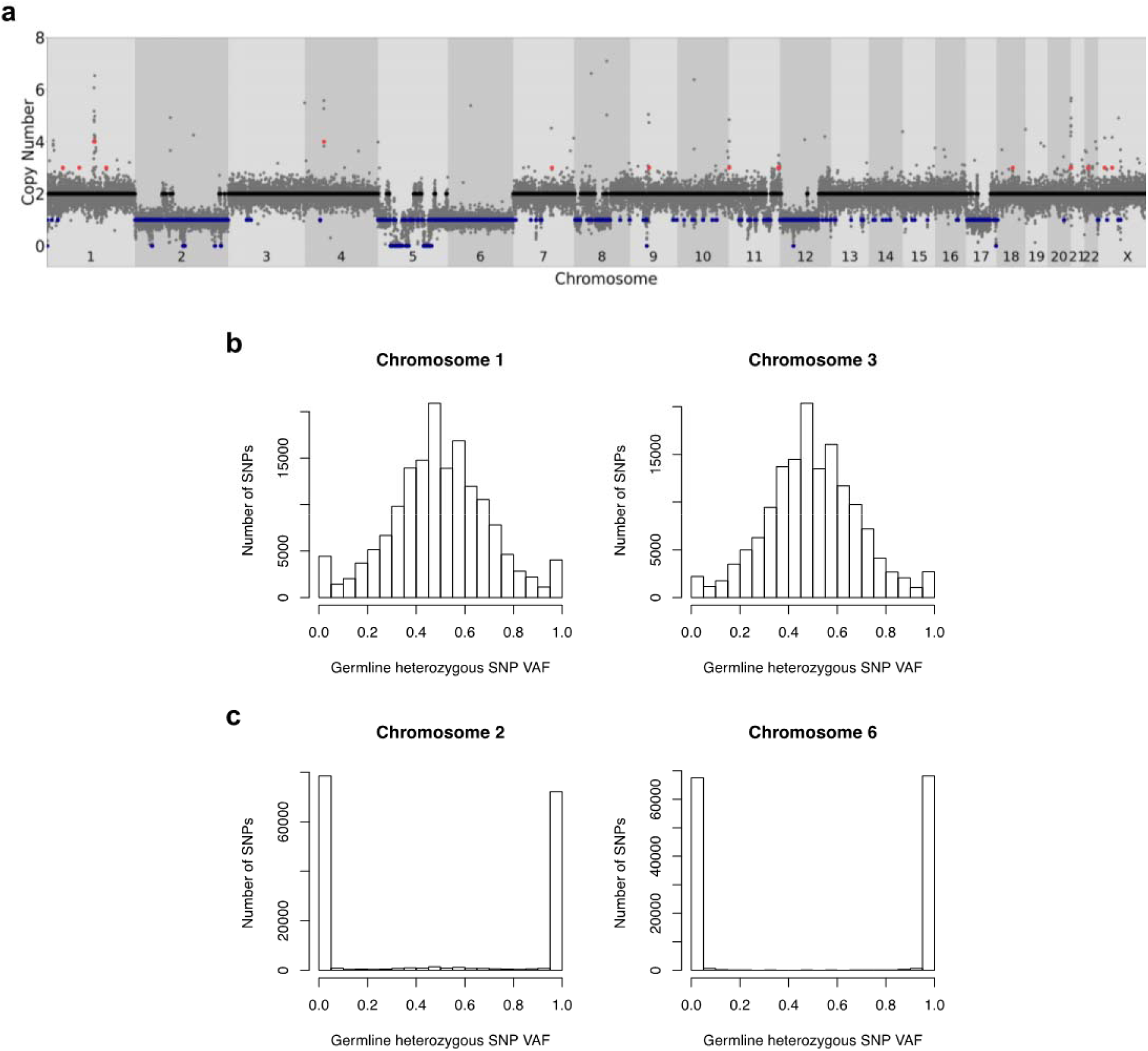
Possible failed PTA amplification. **a.** Neuron B from subject 5823 shows single copy loss over the majority of chromosomes 2, 5, 6, 12 and 17. **b.** Variant allele fractions (VAF) for heterozygous germline SNPs on chromosomes 1 and 3 show the expected VAF variance for successfully amplified chromosomes. **c.** Same as (b) for chromosomes 2 and 6, which show a loss over the majority of each chromosome. VAF values at 0 and 1 are consistent with the complete loss of a single haplotype, ruling out the possibility that both alleles were present and amplified but to a lower level than other chromosomes. However, whether the single neuron truly contained a single copy loss or if the apparent loss resulted from complete amplification failure of one haplotype cannot be determined.

**Supplementary Figure 3:**
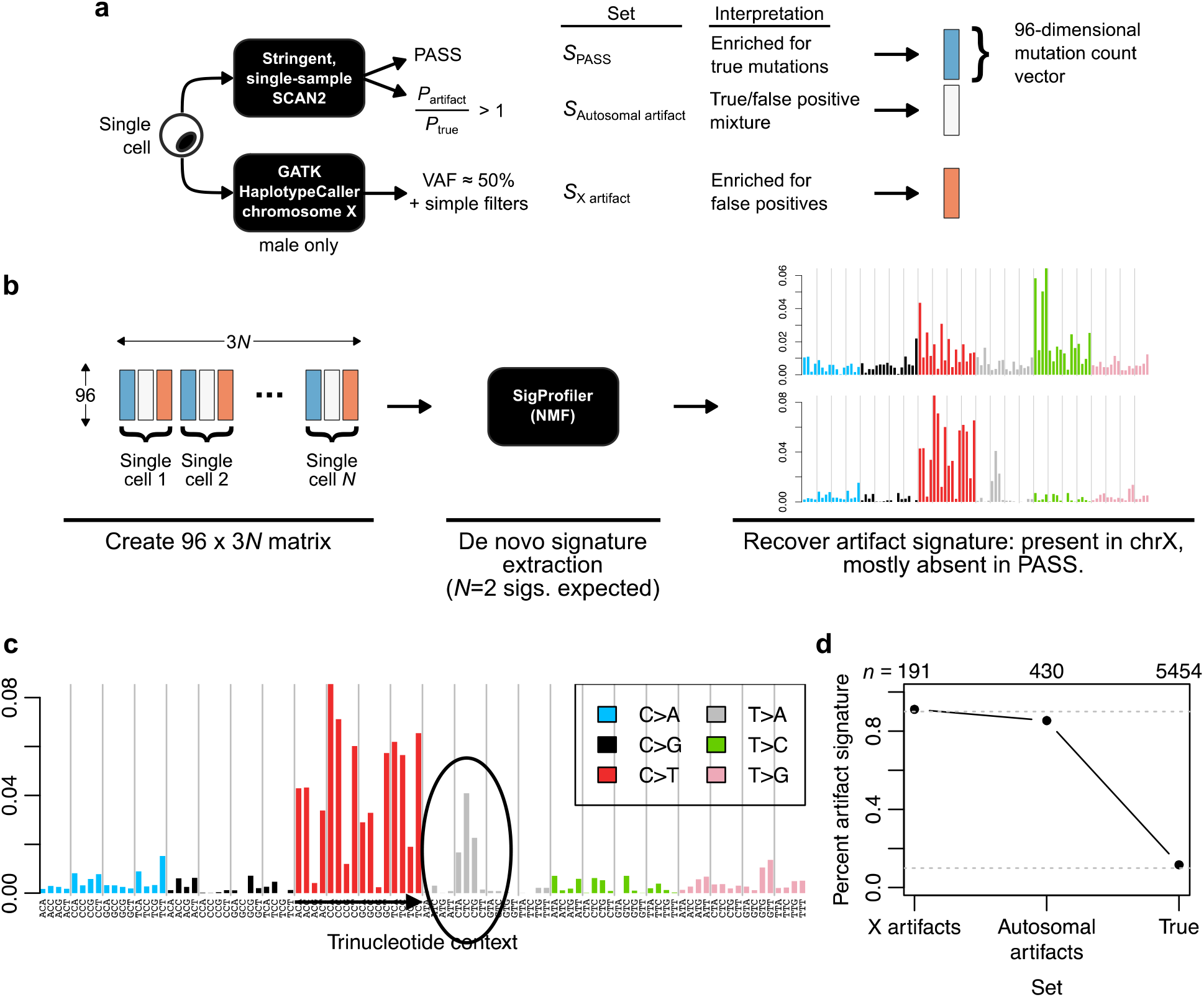
The universal PTA artifact signature. **a.** 3 sets of SNVs and likely artifacts were constructed for each male single cell. PASS autosomal SNVs using stringent calling filters are highly depleted for artifacts while rejected candidate SNVs with *P*_artifact_/*P*_true_ > 1 (see ref. 14 for information on the models corresponding to these *P*-values) or chromosome X sites in the non-pseudoautosomal regions with ~50% VAF in male samples are highly enriched for early, high-VAF PTA artifacts. **b.** An SBS96 mutation count matrix is constructed for de novo signature extraction using 3 separate entries for each male single cell (not shown: female cells are also used but have no X chromosome component). *De novo* signature extraction produced *N*=2 signatures corresponding to the known neuronal aging signature^6^ and the universal PTA artifact signature. **c.** The universal PTA artifact signature in more detail. **d.** Percent of SNVs in each set assigned to the artifact signature by *de novo* extraction. Values (top, *n*) indicate the total number of SNVs in each set from the 25 PTA neurons. Dotted lines: 10% and 90%.

**Supplementary Figure 4:**
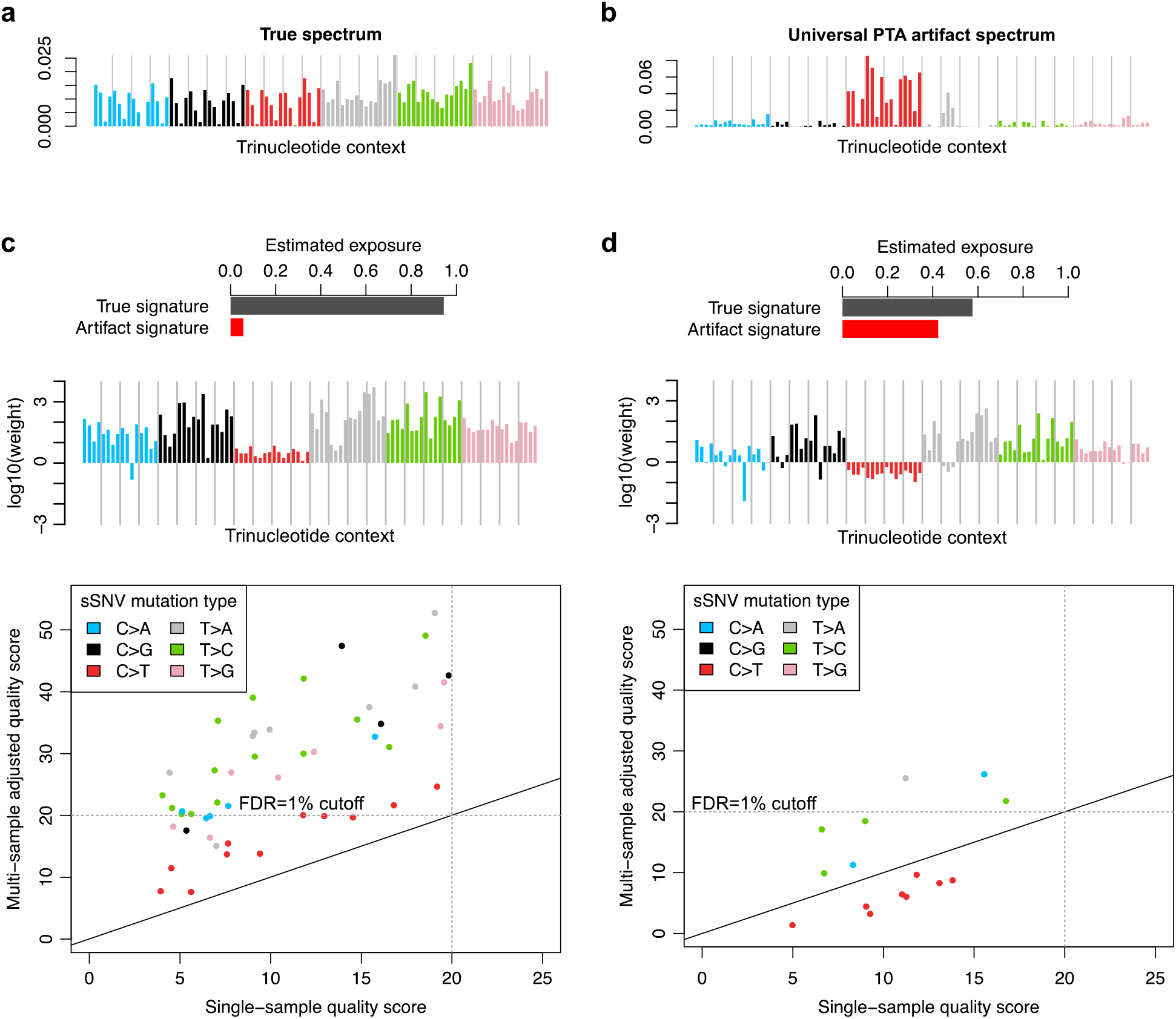
Examples of the multi-sample sSNV approach: weight calculation and quality score adjustment. **a.** True mutation spectrum derived from high confidence calls in simulated data (synthetic diploids, see **Supplementary Figure 6** for a detailed performance comparison). **b.** Universal PTA artifact spectrum (see Methods). **c-d.** Examples of multi-sample adjustment on two single cells (synthetic diploids) with differing artifact burdens. (*Top*) Exposure to the true and artifact mutation signatures derived by least squares fitting; cell-specific exposure to the artifact signature can be interpreted as an estimate of the artifact rate among sSNV candidates. (*Middle*) Log-scaled weights based on estimated artifact exposure, mutation type and trinucleotide context for a specific single cell. (*Bottom*) Adjustment of the FDR heuristic for sSNV candidates from one single cell. Each point represents one sSNV candidate being reconsidered by multi-sample calling. Quality scores are Phred-scaled. Detection threshold of Q=20 corresponds to a target FDR of 0.01. Solid lines, y=x.

**Supplementary Figure 5:**
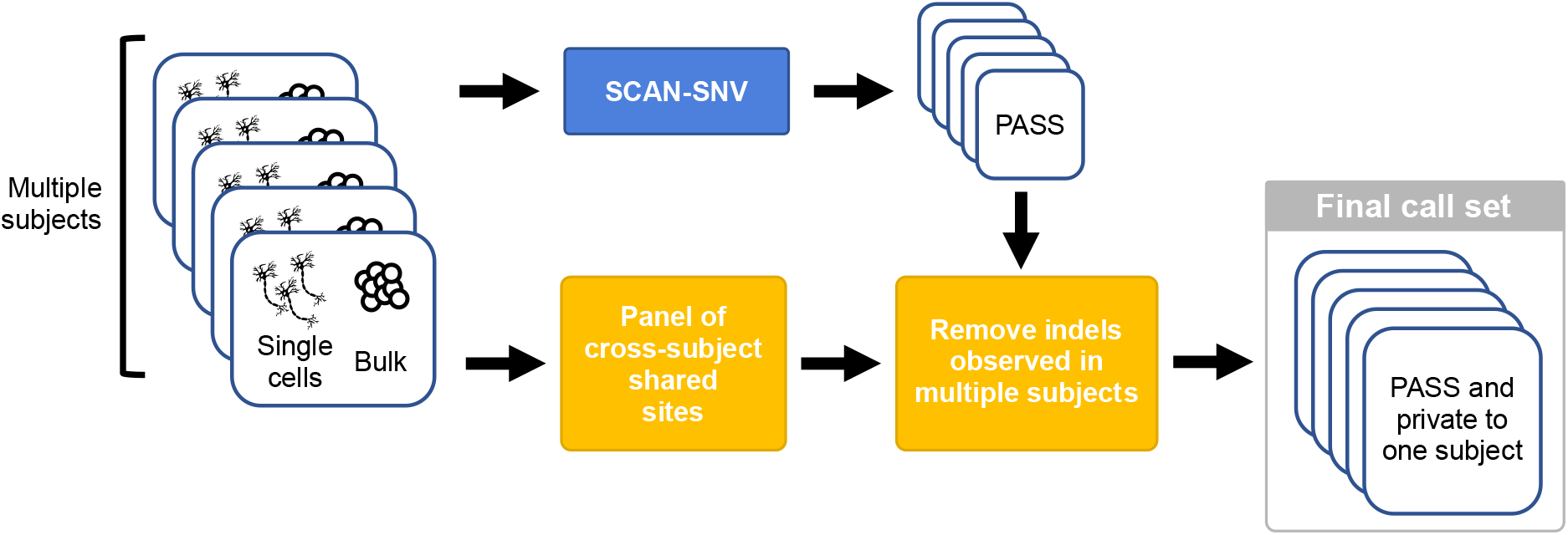
Somatic indel calling strategy. PTA-amplified cells and matching bulk samples from multiple subjects are required for indel calling. Single cells and bulks are each analyzed by a modified SCAN-SNV pipeline. GATK HaplotypeCaller is independently run in joint mode on all single cells and bulks to produce a panel containing reference and alternate read counts across the full cohort. Somatic indels passed by the modified SCAN-SNV pipeline are then removed if reads supporting the indel are observed in single cells from other subjects.

**Supplementary Figure 6:**
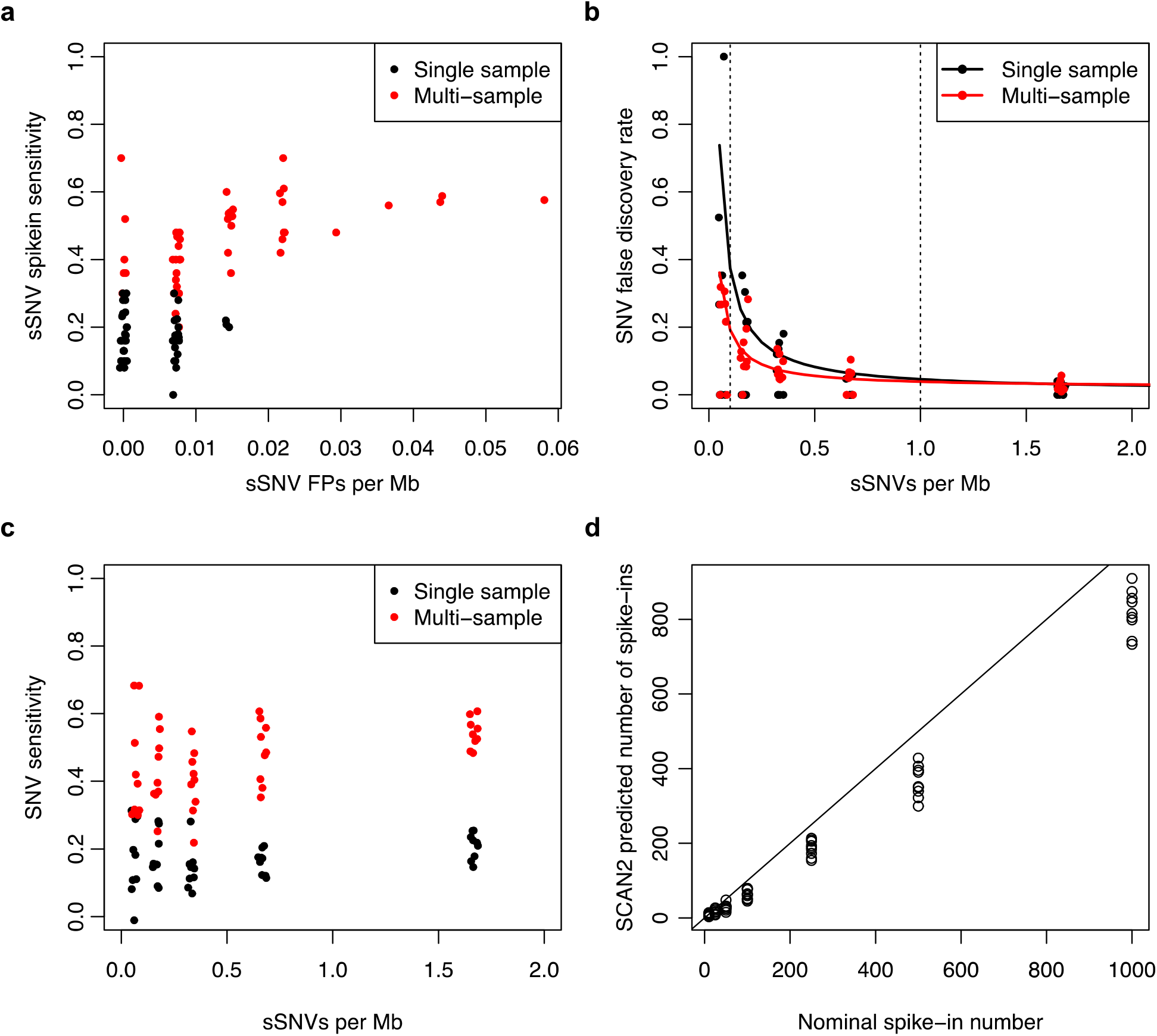
Simulated data to assess sSNV calling performance for single and multi-sample SCAN2. **a.** SNV sensitivity and false positive rate for synthetic diploid simulations with 1-250 spike-ins per simulation. Target FDR=1%, rescue FDR=1%. **b.** SNV sensitivity plotted against mutation burden for simulated SNVs. **c.** False discovery rate plotted against mutation burden for simulated SNVs. Solid lines: linear regression fits to FDR ~ 1/mutations per Mb. Dotted vertical lines: typical range of somatic mutation burdens in healthy single cells. **d.** SCAN2 total sSNV burden estimates for 63 simulations. 9 synthetic diploid simulations were performed for each of the spike-in rates of 10, 25, 50, 100, 250, 500 and 1000 per simulation. Solid line: y=x. x-axes for panels **a-c** are jittered for visibility.

**Supplementary Figure 7:**
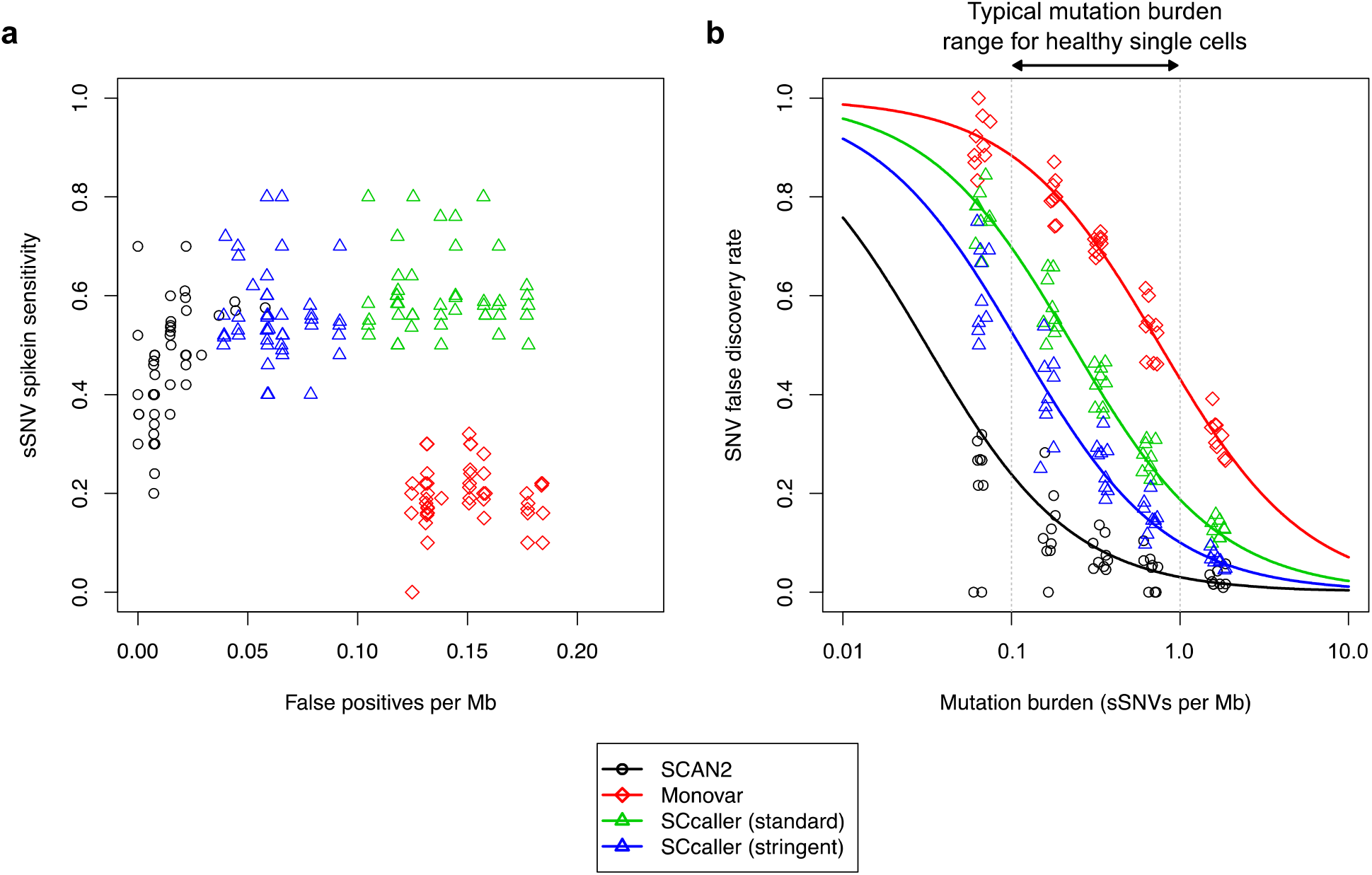
Comparison of SCAN2 to other single-cell SNV genotypers. **a.** Each point represents a single simulated synthetic diploid X chromosome. Sensitivity is the fraction of spike-ins recovered. False positives are SNV calls that were not known spike-ins or endogenous somatic mutations. **b.** False discovery rate vs. the number of spike-ins per megabase. Lines are parameterized by mean sensitivity *S* and false positive rate per megabase *F*: FDR = *F* / (*F* + x*S*). Single cells from non-neoplastic human tissues typically exhibit SNV burdens between 0.1 and 1.0 mutations per Mb (about 250-2500 sSNVs per genome). SCcaller standard uses a calling threshold of *α* = 0.05 while stringent calling uses *α* = 0.01.

**Supplementary Figure 8:**
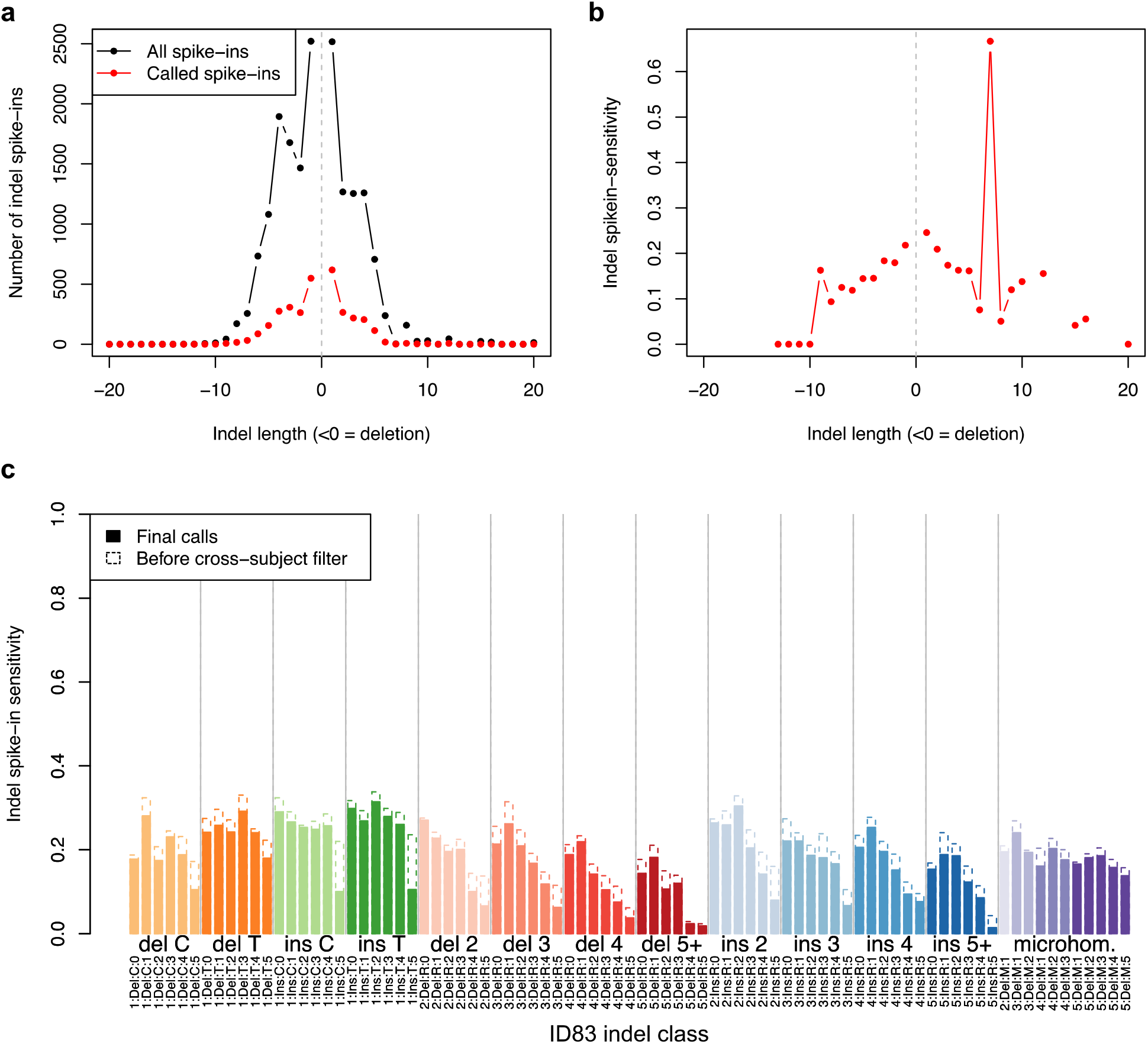
SCAN2 sensitivity on simulated indels. **a.** Length distribution of all simulated spike-in indels (black) and recovered indels (red). **b.** Spike-in indel sensitivity by length. **c.** Sensitivity for indel detection stratified by ID83 indel class. Dotted outlines: sensitivity before applying cross-subject filtration.

**Supplementary Figure 9:**
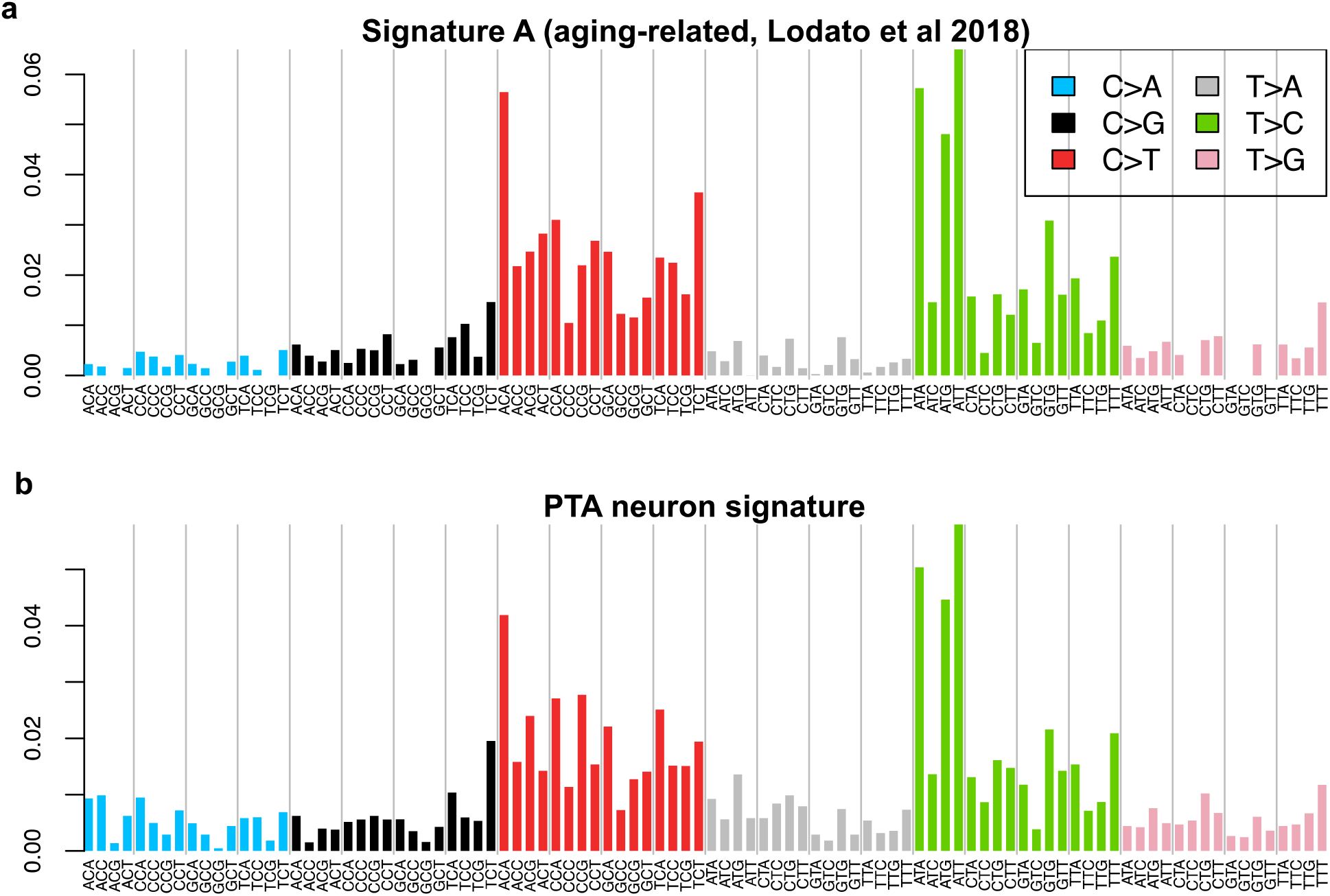
PTA confirms the age-related sSNV signature in human neurons. **a.** Aging-associated signature derived from MDA-amplified neurons (ref. 6)**. b.** Mutation signature produced by single-sample SCAN2 on PTA-amplified human neurons. Multi-sample SCAN2 is not appropriate for mutation signature discovery because it is biased against mutations from signature components with high representation in the universal PTA artifact signature. The PTA neuronal signature is highly similar to Signature A (cosine similarity=0.966), confirming the previously reported signature.

**Supplementary Figure 10:**
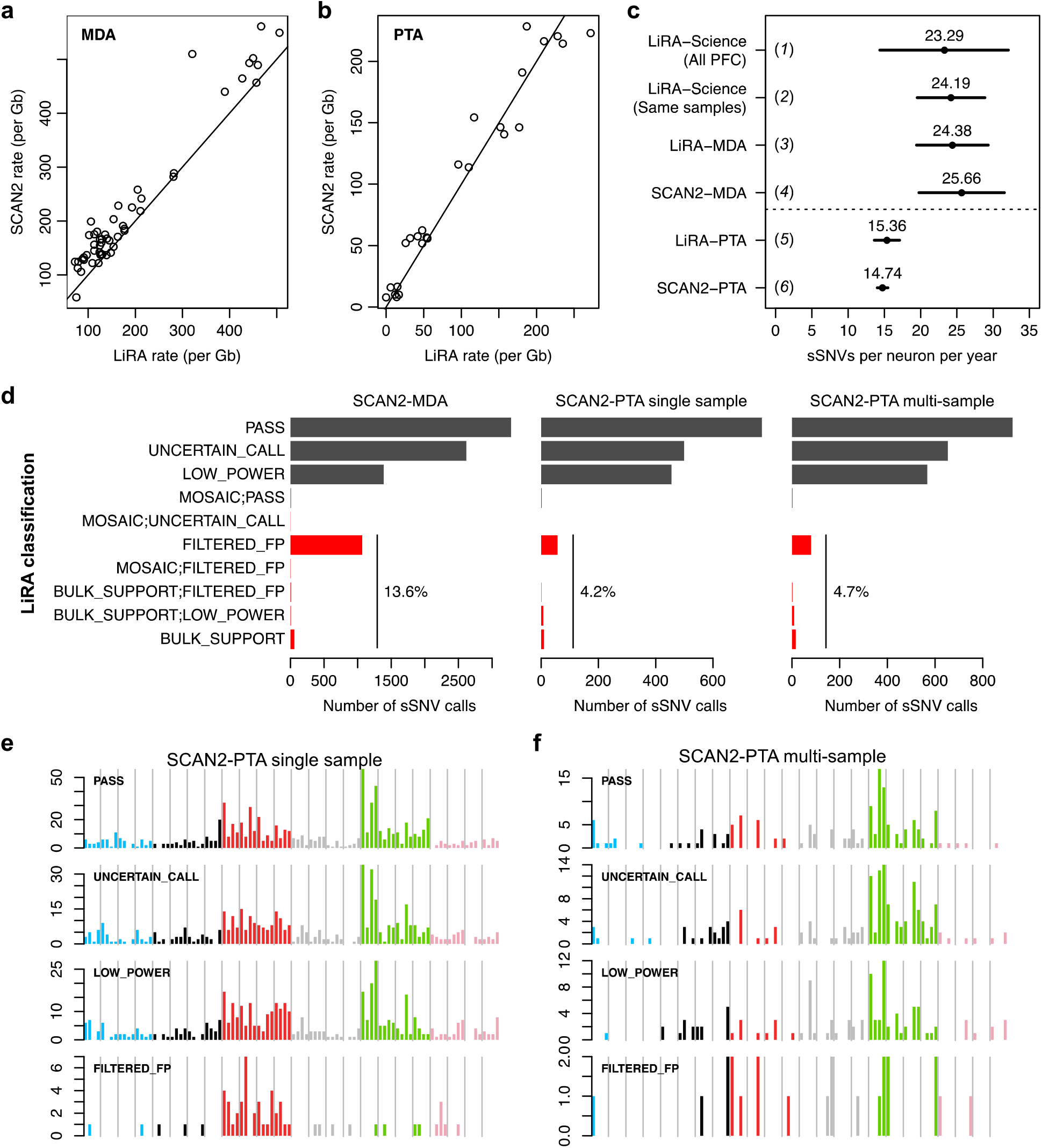
Comparison of SCAN2 and LiRA on human neurons. Single human neurons were previously analyzed by LiRA^20^, a specific but lower sensitivity approach for calling somatic SNVs. **a-b.** SCAN2 and LiRA extrapolations for the total (not called) sSNV burden per diploid Gb of human sequence from MDA-(**a**) and PTA-amplified (**b**) single neurons. Solid lines: y=x. **c.** Linear regression estimates for the number of sSNVs accumulated per neuron per year from several sources and analyses. Horizontal bars represent 95% C.I.s. (*1*) LiRA rates taken from ref. 6, which used a larger set of 91 MDA-amplified PFC neurons; (*2*) LiRA rates taken from ref. 6 using the same set of 51 MDA-amplified PFC neurons; (*3*) rerun of LiRA on 51 MDA-amplified neurons using the same input provided to SCAN2; (*4*) SCAN2 on 51 MDA-amplified neurons; (*5*) LiRA on 25 PTA-amplified neurons; (*6*) SCAN2 on 25 PTA-amplified neurons. **d.** LiRA classification of SCAN2 calls where reads linked to nearby germline heterozygous SNPs are available (black: likely true sSNVs, red: possible false positives). PASS is the highest quality LiRA class. UNCERTAIN and LOW_POWER indicate lack of linking reads to make a confident call, but no evidence of artifactual status is detected. All other classes (red) are interpreted as false positives. Percentages show the fraction of all false positive classes among SCAN2 calls. **e-f.** Raw mutation spectra for single-(**e**) and multi-sample (**f**) SCAN2 calls stratified by LiRA classification. The similarities between PASS and the two lower quality UNCERTAIN_CALL and LOW_POWER classes suggest that the majority of UNCERTAIN_CALL and LOW_POWER SCAN2 calls are true mutations. Confident false positives (FILTERED_FPs) possess a C>T dominated signature with lack of C>Ts at CpGs.

**Supplementary Figure 11:**
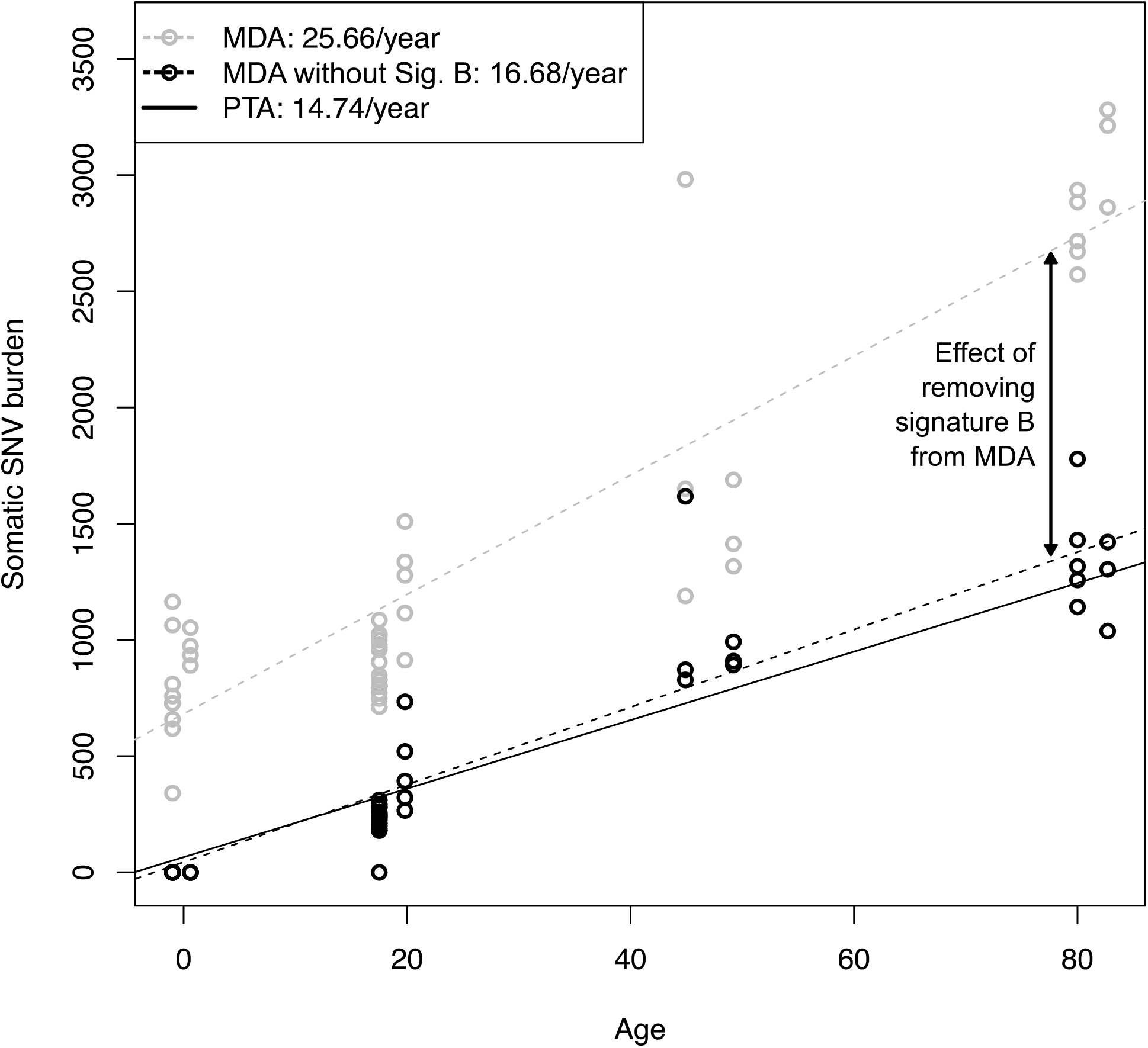
Removal of Signature B from MDA neurons closely matches PTA-derived mutation rates. Total SCAN2-called somatic SNV mutation burdens from MDA neurons before Signature B removal (grey circles) and after Signature B removal (black circles). Trend lines: MDA accumulation rate (dotted grey), MDA accumulation rate after Signature B removal (dotted black), PTA accumulation rate (solid black).

**Supplementary Figure 12:**
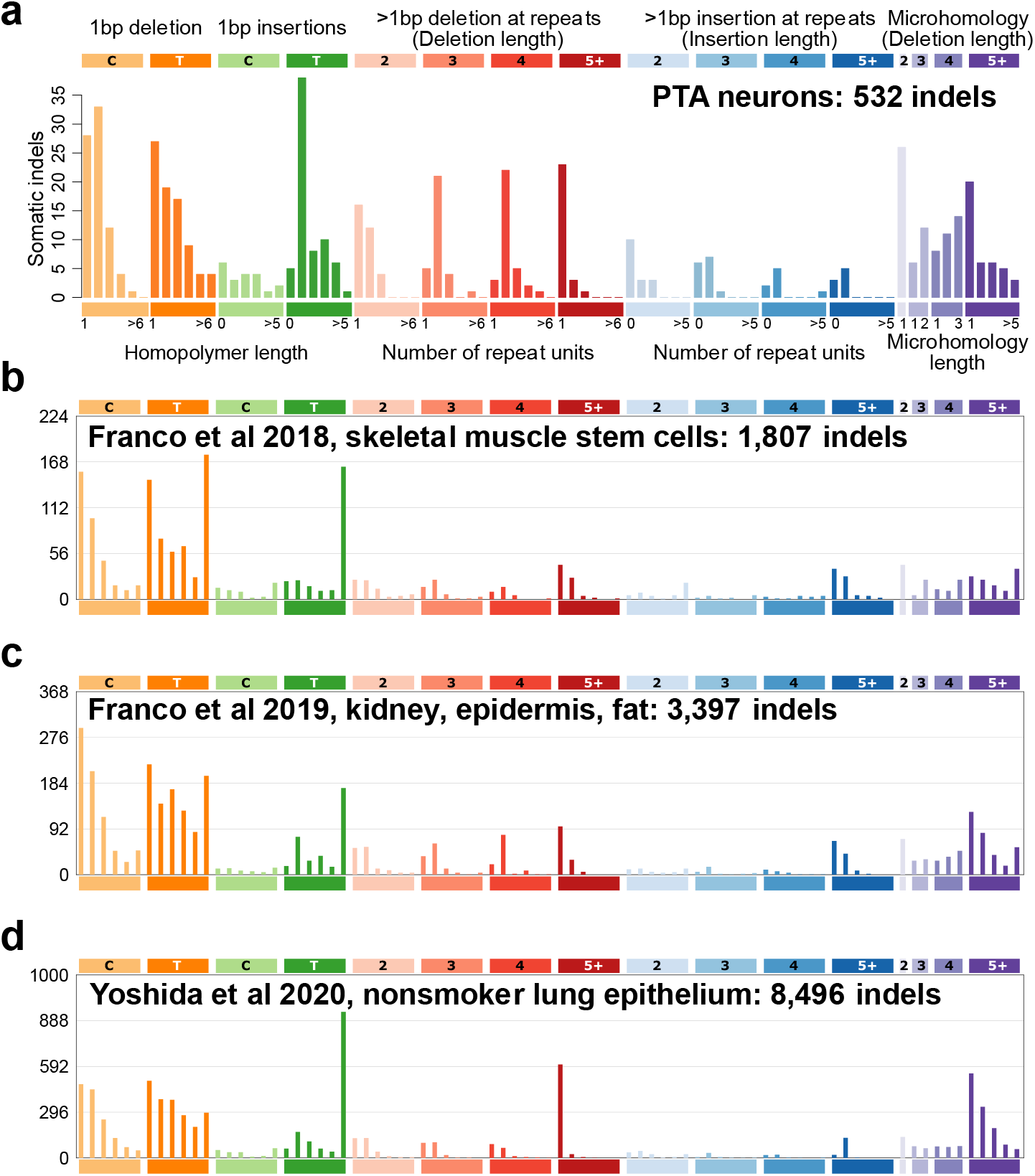
Somatic indel signatures compiled from other publications. **a.** 532 indels from PTA neurons from this study, same as **Figure 3f**. **b.** Clonally expanded single skeletal muscle stem cells. **c.** Clonally expanded single kidney, epidermis and fat cells. Excludes hypermutated kidney cells (designated KT2 in the original study). **d.** Clonally expanded bronchial epithelial cells from children and never-smokers.

**Supplementary Figure 13:**
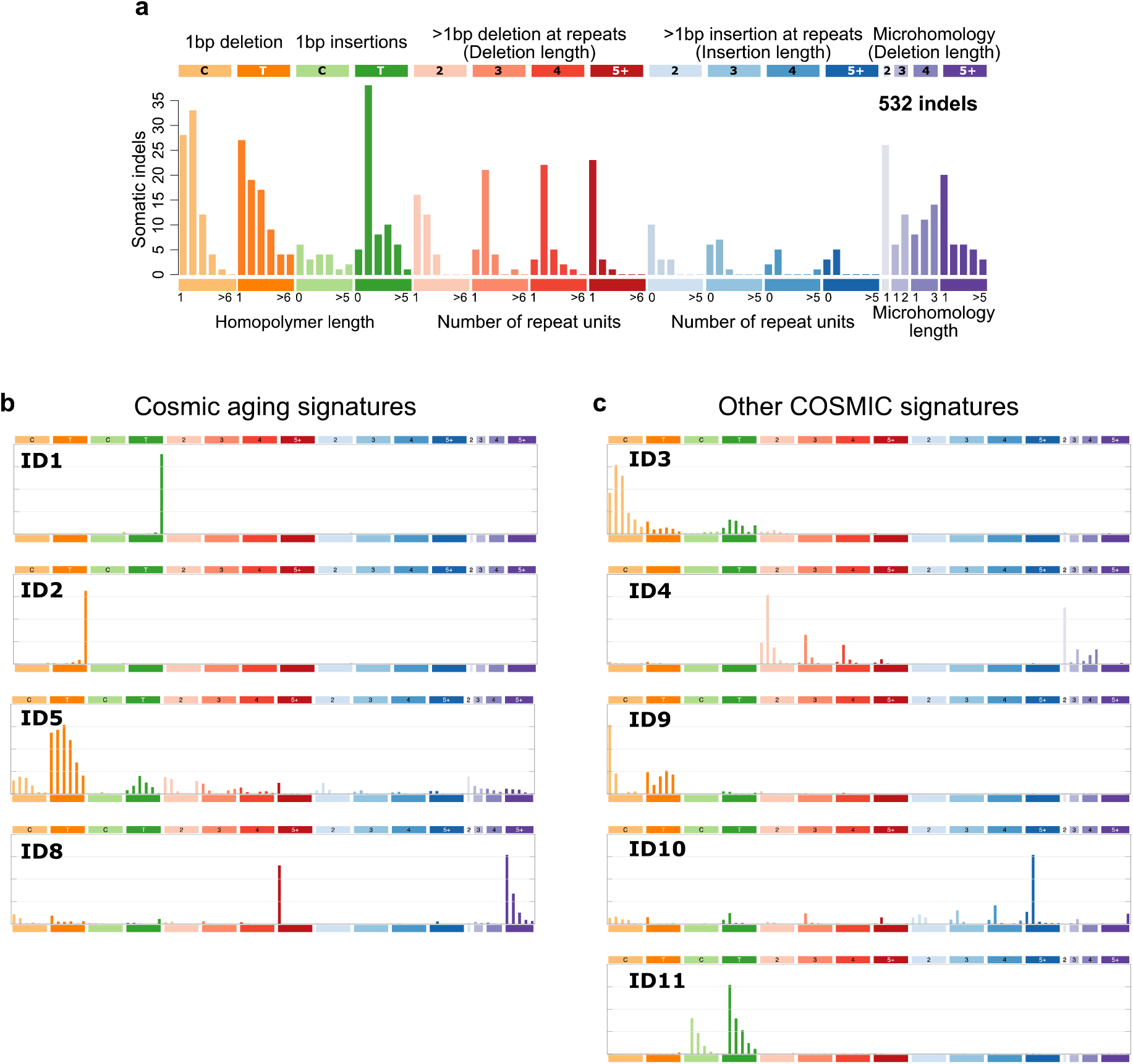
COSMIC indel signatures. **a.** Spectrum of indels from PTA neurons (same as **Figure 3f**). **b.** COSMIC signatures with clock-like or age-associated annotations. **c.** Non-aging COSMIC signatures found in single neurons.

**Supplementary Table 1:** Individuals sequenced in this study. Individuals from four age groups, ranging from infants to the elderly, were analyzed in this study. MDA and PTA columns refer to the number of PFC neurons amplified by each method and sequenced to high coverage.

| Donor ID | Age | Sex | MDA | PTA |
| --- | --- | --- | --- | --- |
| **Infant** |  |  |  |  |
| 1278 | 0.4 | M | 9 | 3 |
| 5817 | 0.6 | M | 4 | 3 |
| **Adolescent** |  |  |  |  |
| 1465 | 17.5 | M | 18 | 4 |
| 5559 | 19.8 | F | 5 | 3 |
| **Adult** |  |  |  |  |
| 5087 | 44.9 | M | 4 | 3 |
| 936 | 49.2 | F | 3 | 3 |
| **Aged** |  |  |  |  |
| 5657 | 82 | M | 5 | 3 |
| 5823 | 82.7 | F | 3 | 3 |

**Supplementary Table 2:** Samples analyzed in this study. List of all samples used in this study. For single cell samples, the method of genome amplification is listed (MDA or PTA); samples with amplification “none” are bulk controls. Callable bp indicates the number of base pairs in the human genome which passed basic depth criteria for analysis (>5 in the single cell, >10 in the matched bulk).

| Subject | Sample | Amp. | Age | Sex | Callable bp | MAPD |
| --- | --- | --- | --- | --- | --- | --- |
| 1278 | 1278_ct_p1E3 | MDA | 0.4 | M | 2377463116 | 0.582 |
| 1278 | 1278_ct_p1E6 | MDA | 0.4 | M | 2254957388 | 0.767 |
| 1278 | 1278_ct_p1G9 | MDA | 0.4 | M | 2310472262 | 0.717 |
| 1278 | 1278_ct_p2B9 | MDA | 0.4 | M | 2294524648 | 0.708 |
| 1278 | 1278_ct_p2C7 | MDA | 0.4 | M | 2351946883 | 0.727 |
| 1278 | 1278_ct_p2E4 | MDA | 0.4 | M | 2277857833 | 0.744 |
| 1278 | 1278_ct_p2E6 | MDA | 0.4 | M | 2315010769 | 0.73 |
| 1278 | 1278_ct_p2F5 | MDA | 0.4 | M | 2298927433 | 0.71 |
| 1278 | 1278_ct_p2G5 | MDA | 0.4 | M | 2285597264 | 0.722 |
| 1278 | 1278BA9-A | PTA | 0.4 | M | 2559227873 | 0.188 |
| 1278 | 1278BA9-B | PTA | 0.4 | M | 2564412679 | 0.187 |
| 1278 | 1278BA9-C | PTA | 0.4 | M | 2570780791 | 0.186 |
| 1278 | 1278_heart_bulk | none | 0.4 | M | NA |  |
| 5817 | 5817_ct_p1H10 | MDA | 0.6 | M | 2218940766 | 0.827 |
| 5817 | 5817_ct_p1H2 | MDA | 0.6 | M | 2264476042 | 0.754 |
| 5817 | 5817_ct_p1H5 | MDA | 0.6 | M | 2280282603 | 0.753 |
| 5817 | 5817_ct_p2H6 | MDA | 0.6 | M | 2241619365 | 0.768 |
| 5817 | 5817PFC-A | PTA | 0.6 | M | 2458549871 | 0.232 |
| 5817 | 5817PFC-B | PTA | 0.6 | M | 2394321853 | 0.226 |
| 5817 | 5817PFC-C | PTA | 0.6 | M | 2432653880 | 0.214 |
| 5817 | 5817_liver_bulk | none | 0.6 | M | NA |  |
| 1465 | 1465-cortex_1-neuron_MDA_12 | MDA | 17.5 | M | 2368914601 | 0.576 |
| 1465 | 1465-cortex_1-neuron_MDA_18 | MDA | 17.5 | M | 2328801902 | 0.569 |
| 1465 | 1465-cortex_1-neuron_MDA_20 | MDA | 17.5 | M | 2343394691 | 0.549 |
| 1465 | 1465-cortex_1-neuron_MDA_24 | MDA | 17.5 | M | 2250564147 | 0.63 |
| 1465 | 1465-cortex_1-neuron_MDA_25 | MDA | 17.5 | M | 2317117886 | 0.574 |
| 1465 | 1465-cortex_1-neuron_MDA_2_WGSb | MDA | 17.5 | M | 2278772444 | 0.553 |
| 1465 | 1465-cortex_1-neuron_MDA_30 | MDA | 17.5 | M | 2278027817 | 0.607 |
| 1465 | 1465-cortex_1-neuron_MDA_39 | MDA | 17.5 | M | 2281108217 | 0.61 |
| 1465 | 1465-cortex_1-neuron_MDA_3_WGSb | MDA | 17.5 | M | 2246826723 | 0.58 |
| 1465 | 1465-cortex_1-neuron_MDA_43 | MDA | 17.5 | M | 2311323225 | 0.545 |
| 1465 | 1465-cortex_1-neuron_MDA_46 | MDA | 17.5 | M | 2329270490 | 0.565 |
| 1465 | 1465-cortex_1-neuron_MDA_47 | MDA | 17.5 | M | 2276931799 | 0.57 |
| 1465 | 1465-cortex_1-neuron_MDA_5 | MDA | 17.5 | M | 2283876392 | 0.579 |
| 1465 | 1465-cortex_1-neuron_MDA_51_WGSb | MDA | 17.5 | M | 2220441876 | 0.602 |
| 1465 | 1465-cortex_1-neuron_MDA_6_WGSb | MDA | 17.5 | M | 2248579628 | 0.561 |
| 1465 | 1465-cortex_1-neuron_MDA_8 | MDA | 17.5 | M | 2319210026 | 0.628 |
| 1465 | 1465_ct_8p2h8 | MDA | 17.5 | M | 2396680438 | 0.548 |
| 1465 | 1465_ctx_p2g8 | MDA | 17.5 | M | 2346370110 | 0.58 |
| 1465 | 1465BA9-A | PTA | 17.5 | M | 2502902455 | 0.207 |
| 1465 | 1465BA9-B | PTA | 17.5 | M | 2379712047 | 0.28 |
| 1465 | 1465BA9-C | PTA | 17.5 | M | 2490365389 | 0.232 |
| 1465 | 1465BA9-D | PTA | 17.5 | M | 2385020412 | 0.272 |
| 1465 | 1465-cortex_BulkDNA_WGSb | none | 17.5 | M | NA |  |
| 5559 | 5559-pfc1C4 | MDA | 19.8 | F | 2335438836 | 0.696 |
| 5559 | 5559-pfc1C7 | MDA | 19.8 | F | 2219634045 | 0.819 |
| 5559 | 5559-pfc1E2 | MDA | 19.8 | F | 2243450288 | 0.861 |
| 5559 | 5559-pfc1H2 | MDA | 19.8 | F | 2177525787 | 0.815 |
| 5559 | 5559-pfc2A3 | MDA | 19.8 | F | 2380863121 | 0.701 |
| 5559 | 5559PFC-A | PTA | 19.8 | F | 2506510681 | 0.21 |
| 5559 | 5559PFC-B | PTA | 19.8 | F | 2474056125 | 0.206 |
| 5559 | 5559PFC-C | PTA | 19.8 | F | 2532204421 | 0.193 |
| 5559 | 5559-bulk | none | 19.8 | F | NA |  |
| 5087 | 5087pfc-Lp1C5 | MDA | 44.9 | M | 1956709694 | 1.105 |
| 5087 | 5087pfc-Rp1G4 | MDA | 44.9 | M | 2045937656 | 1.027 |
| 5087 | 5087pfc-Rp3C5 | MDA | 44.9 | M | 851472780 | 1.758 |
| 5087 | 5087pfc-Rp3F4 | MDA | 44.9 | M | 2219472588 | 0.848 |
| 5087 | 5087PFC-A | PTA | 44.9 | M | 2526638419 | 0.192 |
| 5087 | 5087PFC-B | PTA | 44.9 | M | 2529648486 | 0.199 |
| 5087 | 5087PFC-C | PTA | 44.9 | M | 2496175648 | 0.194 |
| 5087 | 5087-hrt-1b1 | none | 44.9 | M | NA |  |
| 936 | 936_20141001-pfc-1cp1G11_20170221-WGS | MDA | 49.2 | F | 2069054494 | 0.95 |
| 936 | 936_20141001-pfc-1cp1H9_20170221-WGS | MDA | 49.2 | F | 2239568087 | 0.85 |
| 936 | 936_20141001-pfc-1cp2F6_20170221-WGS | MDA | 49.2 | F | 1937342351 | 1.036 |
| 936 | 936PFC-A | PTA | 49.2 | F | 2458178187 | 0.189 |
| 936 | 936PFC-B | PTA | 49.2 | F | 2498078321 | 0.186 |
| 936 | 936PFC-C | PTA | 49.2 | F | 2448162449 | 0.183 |
| 936 | 936-hrt-1b1_20170221-WGS | none | 49.2 | F | NA |  |
| 5657 | 5657-pfc1D2 | MDA | 82 | M | 1970196085 | 1.076 |
| 5657 | 5657-pfc1E11 | MDA | 82 | M | 2358110993 | 0.771 |
| 5657 | 5657-pfc2A6 | MDA | 82 | M | 2379723437 | 0.74 |
| 5657 | 5657-pfc2F1 | MDA | 82 | M | 2397069500 | 0.728 |
| 5657 | 5657-pfc2G9 | MDA | 82 | M | 2405157253 | 0.748 |
| 5657 | 5657PFC-A | PTA | 82 | M | 2477582773 | 0.191 |
| 5657 | 5657PFC-B | PTA | 82 | M | 2531288033 | 0.187 |
| 5657 | 5657PFC-C | PTA | 82 | M | 2467010743 | 0.185 |
| 5657 | 5657-bulk | none | 82 | M | NA |  |
| 5823 | 5823_20160824-pfc-1cp1F11_20170221-WGS | MDA | 82.7 | F | 2096166634 | 1.078 |
| 5823 | 5823_20160824-pfc-1cp2E1_20170221-WGS | MDA | 82.7 | F | 1878096891 | 1.143 |
| 5823 | 5823_20160824-pfc-1cp2G5_20170221-WGS | MDA | 82.7 | F | 1901575408 | 1.062 |
| 5823 | 5823PFC-A | PTA | 82.7 | F | 2435263867 | 0.194 |
| 5823 | 5823PFC-B | PTA | 82.7 | F | 2399384951 | 0.215 |
| 5823 | 5823PFC-C | PTA | 82.7 | F | 2494418160 | 0.19 |
| 5823 | 5823-tempmusc-1b1_20170221-WGS | none | 82.7 | F | NA |  |

